# The H3K27me3 reader UAD-2 recruits a TAF-12-containing transcription condensates to initiate piRNA expression within heterochromatic clusters

**DOI:** 10.1101/2025.04.11.648483

**Authors:** Xinya Huang, Yan Kuang, Meili Li, Minjie Hong, Demin Xu, Jiewei Cheng, Tongtong Huang, Xianlei Sun, Wenkai Wang, Ying Zhou, Xuezhu Feng, Xiangyang Chen, Chengming Zhu, Shouhong Guang

## Abstract

Metazoans utilize the small RNA pathway to regulate gene expression and maintain genome integrity. This pathway directs histone H3 lysine 9 tri-methylation (H3K9me3) or histone H3 lysine 27 tri-methylation (H3K27me3) at target loci to induce transcriptional gene silencing. Interestingly, some small RNAs are generated from genomic loci enriched in H3K9me3 or H3K27me3. However, the transcription mechanism of small RNA precursors from these heterochromatic regions remains unclear. In *C. elegans*, piRNAs originate from two genomic clusters enriched with H3K27me3 marks, which recruit the H3K27me3 reader UAD-2 and the upstream sequence transcription complex (USTC). Here, we demonstrate that piRNA transcription in *C. elegans* relies on TAF-12, a subunit of the basal transcription factor IID (TFIID). Depletion of TAF-12 reduces the production of both piRNA precursors and mature piRNAs. TAF-12 interacts with UAD-2 and facilitates piRNA focus formation in germ cell nuclei. We further show that TAF-12 triggers piRNA transcription by recruiting the RNA polymerase II subunit RPB-5, the Mediator complex subunit MDT-8, and the general transcription factors GTF-2F2 and GTF-2H2C. Thus, piRNA transcription within heterochromatic regions depends on the collaboration between histone modification readers, piRNA-specific transcription factors, and core transcription machinery.

## Introduction

PIWI-interacting RNAs (piRNAs) are functionally conserved in metazoans, while vary in length and sequence among different species. piRNAs associate with Argonaute proteins of the PIWI clade to safeguard genome integrity by suppressing foreign genetic elements at both transcriptional and post-transcriptional levels ^1–7^. Besides, the piRNA pathway is involved in fertility, sex determination, viral defense, and transgenerational inheritance ^8–16^. In *C. elegans*, piRNAs are also named 21U-RNAs, given that they are predominantly 21 nucleotides in length and bear a 5*’* terminal uridine residue ^2,17–20^. There are two types of piRNA genes. Type I piRNA genes are located in two piRNA clusters on chromosome IV and possess the Ruby motif upstream of their transcription start sites. In contrast, type II piRNAs are much less abundant, often transcribed bidirectionally from the transcription start sites of coding genes, and likely lack the Ruby motif ^19–21^.

*C. elegans* piRNA genes are individually transcribed by RNA polymerase II (Pol II). The transcription of piRNA precursors relies on the upstream sequence transcription complex (USTC), which is composed of PRDE-1, SNPC-4, TOFU-4 and TOFU-5 ^22–24^. The USTC complex coats two piRNA clusters by binding to the Ruby motif upstream of type I piRNA gene’s promoters and forms distinct piRNA foci in germ cells. Casein kinase II (CK2) directly phosphorylates the TOFU-4 protein to promote the USTC assembly ^25^. Interestingly, piRNA clusters exhibit signatures of facultative heterochromatin mark histone 3 lysine 27 tri-methylation (H3K27me3) ^26^. H3K27me3 is catalyzed by the Polycomb repressive complex 2 (PRC2), which in *C. elegans* consists of MES-2, MES-3, and MES-6 ^27–29^. Knockdown of MES-2, MES-3, or MES-6 results in dispersed piRNA foci of the USTC complex and reduced piRNA levels ^30^. UAD-2 recognizes and binds to the H3K27me3 mark via its chromodomain, enhancing the USTC complex’s binding to piRNA clusters ^30,31^. UAD-2 accumulates in nuclear piRNA foci and exhibits properties of liquid-liquid phase separation to facilitate the assembly of piRNA transcription machinery ^32^. Additionally, the chromatin remodeling factor ISW-1 recruits the USTC complex to organize the local nucleosome environment upstream of individual piRNA genes and maintains high nucleosome density across piRNA clusters ^30,33^. However, the mechanism by which H3K27me3 promotes piRNA production and how piRNA precursors are transcribed from heterochromatic clusters remain mysterious.

After piRNA transcription initiation, the RNA polymerase II subunit RPB-9 recruits the integrator complex to terminate the transcription of piRNA precursors ^34,35^. These precursors are exported from the nucleus and bound by the PICS/PETISCO complex, which is enriched in the E granule, to stabilize piRNA precursors ^36–40^. piRNA precursors are then cleaved at their 5*’*-terminal by the PUCH complex ^41^. After the removal of the 5*’* m^7^G-cap and the first two nucleotides, piRNA precursors are loaded onto the PIWI protein PRG-1 ^2^. The 3*’*-terminal of piRNA precursors is trimmed by the conserved exonuclease PARN-1 and 2*’*-O-methylated by the RNA methyltransferase HENN1 to generate mature piRNAs ^42–45^. The processing and maturation of piRNAs may occur sequentially in distinct perinuclear germ granules ^40^.

In *Drosophila melanogaster*, most piRNA genes are transcribed from dual-strand piRNA clusters enriched in the heterochromatic H3K9me3 mark ^46^. Rhino, a paralogue of heterochromatin protein-1 (HP1), specifically binds to H3K9me3-enriched chromosomal regions through its chromodomain ^47,48^. The zinc finger protein Kipferl recruits Rhino to guanosine-rich DNA motifs present at piRNA source loci and stabilizes it on chromatin ^49^. Rhino interacts with Deadlock and Cutoff (known as the RDC complex) to promote non-canonical transcription from these loci ^50–53^. The RDC complex further recruits the germline-specific TFIIA-L paralogue Moonshiner, the transcription factor TFIIA-S, and the TATA-box binding protein (TBP)-related factor TRF2 to initiate piRNA transcription ^54^. Interestingly, the H3K27 methyltransferase E(z) guides Rhino to Kipferl-independent piRNA source loci to regulate transposable element expression and piRNA production in *Drosophila* germ cells ^55^, suggesting that the mechanism by which heterochromatin marks promote piRNA transcription is evolutionarily conserved across species.

Pol II initiates transcription as part of the pre-initiation complex (PIC), which consists of Pol II, the Mediator complex, and a set of general transcription factors (GTFs), including TFIIA, TFIIB, TFIID, TFIIE, TFIIF, and TFIIH ^56–58^. Among these GTFs, the TATA binding protein (TBP), a subunit of TFIID, binds directly to core promoters, facilitating promoter melting and transcription initiation. This process is stabilized by TFIIA and TFIIB through direct contacts with TBP. TFIIB further recruits Pol II, TFIIE, TFIIF, and TFIIH. Finally, Mediator binds to GTFs that occupy the enhancer to yield a functional PIC. However, in *C. elegans*, most piRNA source loci are located in facultative heterochromatin regions, suggesting that the initiation mechanism of piRNA transcription may differ from that of protein-coding genes.

Here, we identify that the H3K27me3 reader UAD-2 directly interacts with the TFIID transcription factor TAF-12 to drive piRNA transcription. UAD-2 utilizes its C-terminal domain to bridge TAF-12 and TOFU-4. TAF-12 accumulates in piRNA foci, and its localization is dependent on UAD-2 and the USTC complex. TAF-12 contains a histone-fold domain and a conserved C-terminal domain, both of which are essential for TAF-12’s function in piRNA production. Additionally, the RNA Pol II subunit RPB-5, the Mediator complex subunit MDT-8, the TFIIF subunit GTF-2F2, and the TFIIH subunit GTF-2H2C promote piRNA production by forming a piRNA-specific transcription complex with TAF-12. Overall, our data provide key insights into the transcription mechanism of piRNAs within heterochromatic regions.

## Results

### Identification of TAF-12 interacting with UAD-2 to drive piRNA transcription

To elucidate the molecular mechanism of UAD-2 in piRNA transcription, we performed immunoprecipitation (IP) experiments from whole-worm lysates of *uad-2::gfp::3xflag* animals and identified putative UAD-2-interacting partners through quantitative mass spectrometry (MS). The most enriched protein was TAF-12, a TBP-associated transcription factor (Fig. 1a and Fig. S1a). Using FLAG-tagged TAF-12 as bait, we conversely substantiated the interaction between UAD-2 and TAF-12 via the IP-MS method (Fig. 1b). TAF-12 contains a histone-fold domain and is highly conserved in metazoans as a component of the general transcription factor complex TFIID (Fig. 1c), which plays a central role in RNA polymerase II (Pol II)-dependent transcription initiation ^59–61^. In contrast to *uad-2* mutants, which are viable with developed gonads, *taf-12* mutants are embryonically lethal ^30^. Thus, we utilized the auxin-inducible degron (AID) system to specifically deplete TAF-12 proteins in the germline (Fig. 1d). The knockdown of TAF-12 dramatically reduced UAD-2::GFP fluorescence intensity and abolished piRNA focus formation of UAD-2 (Fig. 1e). This also resulted in the reductions of *uad-2* mRNA and UAD-2::GFP protein levels (Fig. 1f and Fig. S1b). The TAF-12::tagRFP (LG II) rescue experiment confirmed that TAF-12 is essential for the formation of UAD-2 condensates (Fig. 1e).

**Figure 1.**
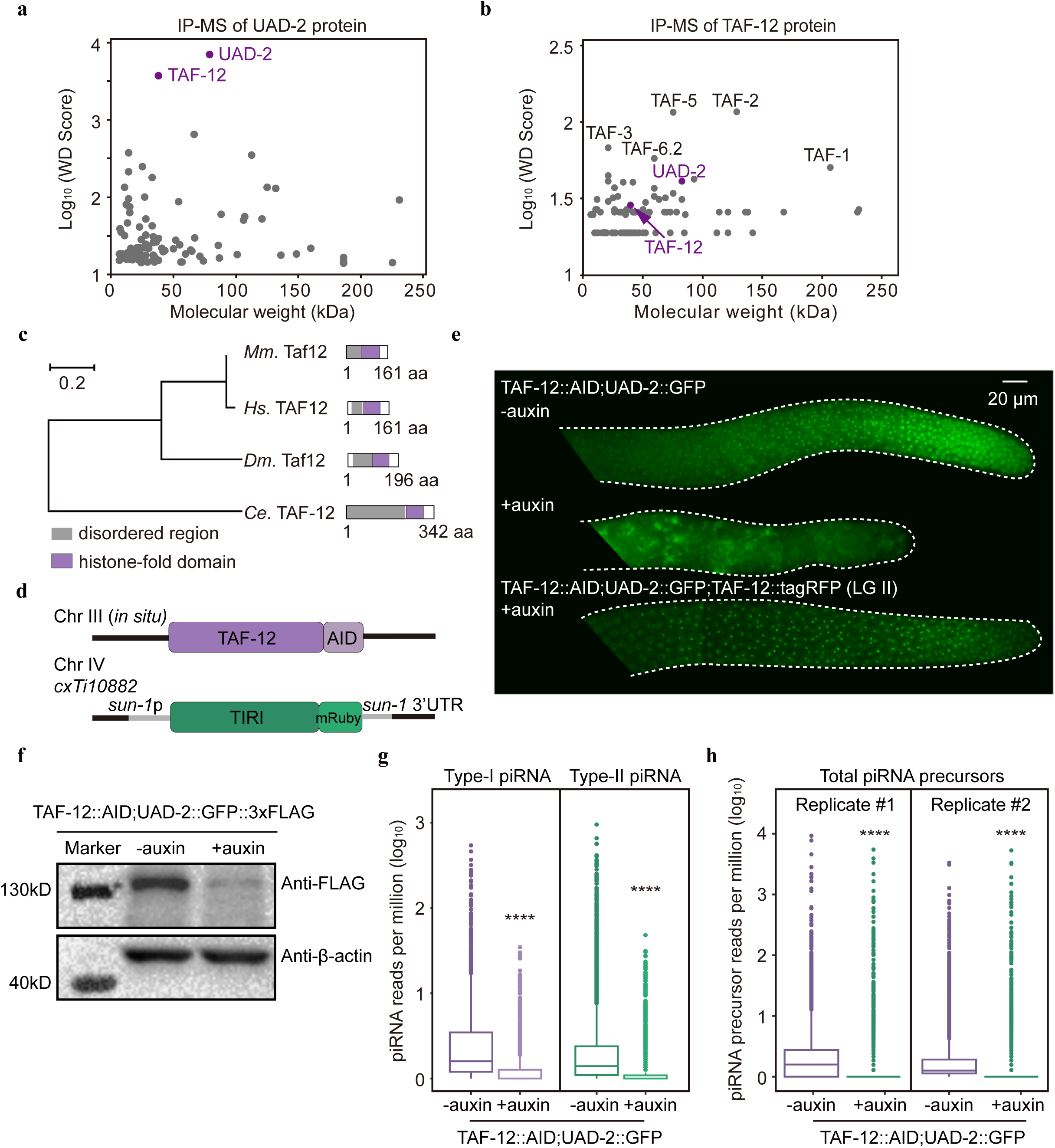
TAF-12 interacts with UAD-2 to drive piRNA transcription. **a-b,** Scatter plots showing (**a**) GFP-tagged UAD-2 and (**b**) 3xFLAG-tagged TAF-12 interaction partners by immunoprecipitation followed by mass spectrometry (IP-MS). Y-axis, Log_10_ (WD score). X-axis, Molecular weight (kDa) of UAD-2 or TAF-12 co-immunoprecipitating proteins. **c,** Phylogenetic tree of TAF-12 in indicated species. TAF-12 encodes a disordered region (grey) and a histone-fold domain (purple). **d,** Schematic of the *taf-12::AID* knock-in allele (Chr III, *in situ*) generated by CRISPR-Cas9 technology in *ieSi38 [sun-1p::TIR1::mRuby::sun-1_3’UTR + Cbr-unc-119(+)]* (Chr IV) background. **e,** Widefield fluorescence microscopy analysis of an adult hermaphrodite expressing UAD-2::GFP and TAF-12::AID with or without auxin treatment, indicating localization and fluorescence intensity of UAD-2::GFP. TAF-12::tagRFP (LG II) rescued UAD-2::GFP focus formation under auxin treatment. Germlines are outlined by white dashed lines. Scale bar, 20 μm. **f,** Western blotting analysis of the expression levels of UAD-2::GFP::3xFLAG and β-actin with or without auxin treatment using anti-FLAG and anti-β-actin antibodies. **g-h,** The relative abundance of (**g**) mature piRNAs from individual type-I and type-II piRNA loci and (**h**) piRNA precursors in young adult hermaphrodites expressing UAD-2::GFP and TAF-12::AID with or without auxin treatment.

In *C. elegans*, PRG-1 is the only functional PIWI clade Argonaute protein and can be loaded with piRNAs to silence transposable elements ^2^. The expression level and perinuclear localization of PRG-1 depend on piRNA accumulation. Loss of piRNA biogenesis factors, such as PRDE-1, KIN-3, and TOFU-6, dramatically reduces the expression level of PRG-1 ^23,25,36^. Thus, we utilized *gfp::prg-1* animals as a piRNA reporter. The knockdown of TAF-12 disrupted the perinuclear localization of PRG-1 and caused a remarkable reduction in PRG-1 protein levels (Fig. S1c-d). Small RNA sequencing further revealed a decrease in both piRNA precursors and mature piRNAs following germline-specific knockdown of TAF-12 (Fig. 1g-h and Fig. S1e). Thus, TAF-12 interacts with UAD-2 and is crucial for piRNA production.

### piRNA focus formation of TAF-12, UAD-2, and the USTC complex is interdependent

Given its interaction with UAD-2, we hypothesized that TAF-12 is enriched at the nuclear piRNA foci in germ cells, similar to UAD-2 and the USTC complex. To test this, we then inserted a *gfp::3xflag* tag to the *taf-12* genomic locus and examined the expression pattern of TAF-12. In embryos, somatic cells, and germ cells, TAF-12 is constitutively expressed and colocalizes with the histone protein HIS-58 (Fig. S2a-c). As expected, TAF-12 colocalizes with UAD-2 at the piRNA foci in germ cells (Fig. 2a).

**Figure 2.**
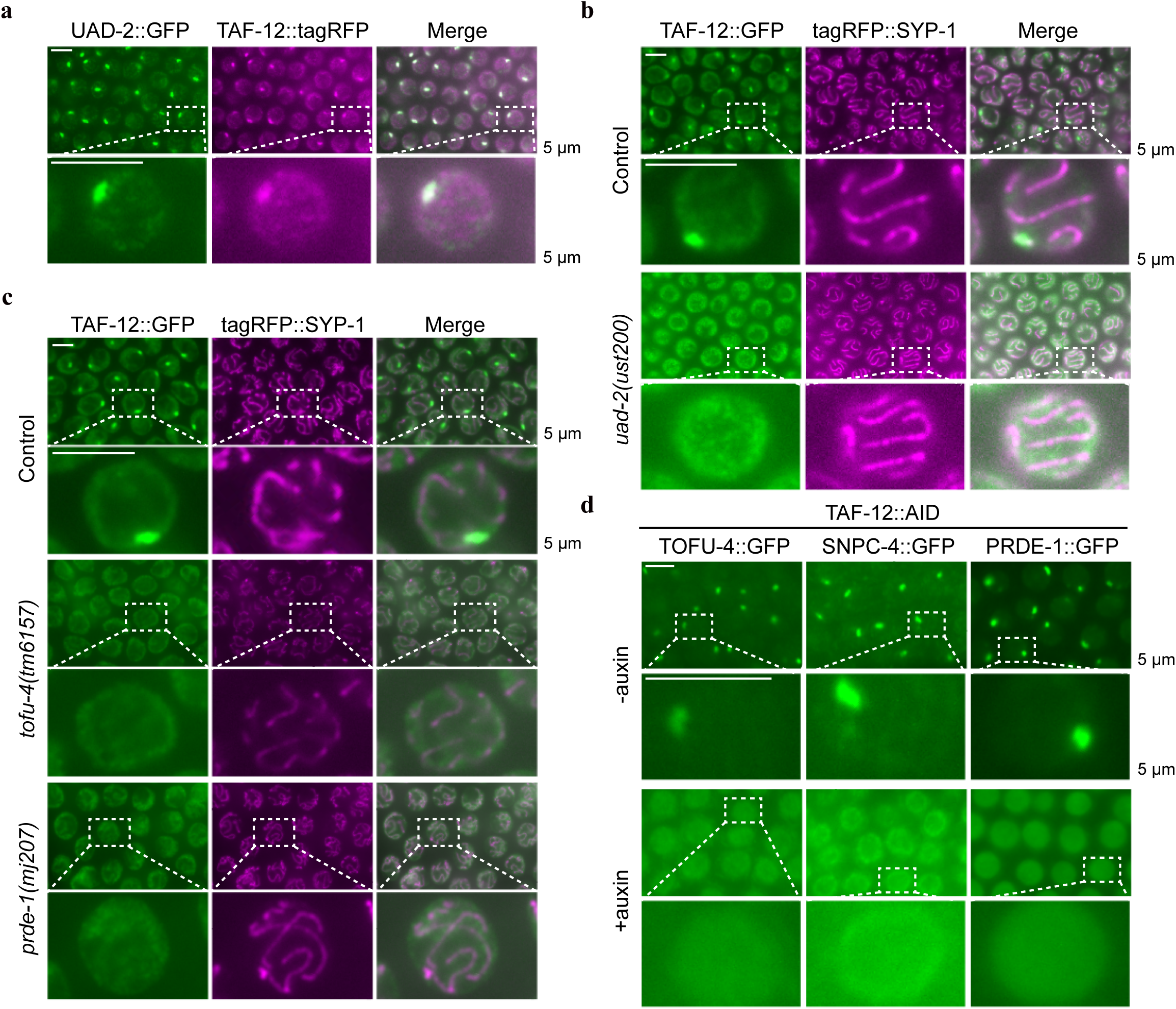
The aggregation of TAF-12, UAD-2, and the USTC complex at piRNA foci is interdependent. **a,** Pachytene germ cells from animals expressing UAD-2::GFP and TAF-12::tagRFP. TAF-12 aggregates in piRNA foci and colocalizes with UAD-2. **b-c,** Pachytene germ cells from animals expressing TAF-12::GFP and tagRFP::SYP-1 in the indicated genetic backgrounds. SYP-1 is a component of the synaptonemal complex and is used as a marker to label the chromosomes. **d,** Fluorescence images of representative pachytene germ cells of the indicated animals expressing TOFU-4::GFP, SNPC-4::GFP or PRDE-1::GFP and TAF-12::AID (with or without auxin treatment). Scale bar, 5 μm.

We used the synaptonemal complex protein SYP-1 as a marker to label chromosomes in pachytene cells ^62^ and examined the cellular localization of TAF-12::GFP in *uad-2* and *ustc* mutants. In the absence of UAD-2, TOFU-4, or PRDE-1, TAF-12 failed to form piRNA foci (Fig. 2b-c). The knockdown of TAF-12 also prevented TOFU-4, SNPC-4, and PRDE-1 from forming piRNA foci (Fig. 2d and Fig. S2d). The protein level of TAF-12 was largely unchanged in *uad-2* and *tofu-4* mutants, but was reduced to approximately 37% in *prde-1* mutants (Fig. S2e-f). Together, these results reveal that the enrichment of TAF-12, UAD-2, and the USTC complex at piRNA foci is mutually dependent.

### UAD-2 binds TAF-12 and the USTC complex through its C-terminal domain

To assess the molecular connections between TAF-12, UAD-2, and the USTC complex, we examined protein-protein interactions among these factors using yeast two-hybrid (Y2H) assays ^22^. Consistent with previous results, TOFU-4 interacts with PRDE-1, and TAF-12 interacts with UAD-2 (Fig. 1a-b, 3a and Fig. S3a) ^22^. Notably, UAD-2 also interacts with TOFU-4 (Fig. 3a and Fig. S3a), which supports with the mutual dependency between UAD-2 and the USTC complex for piRNA focus formation.

**Figure 3.**
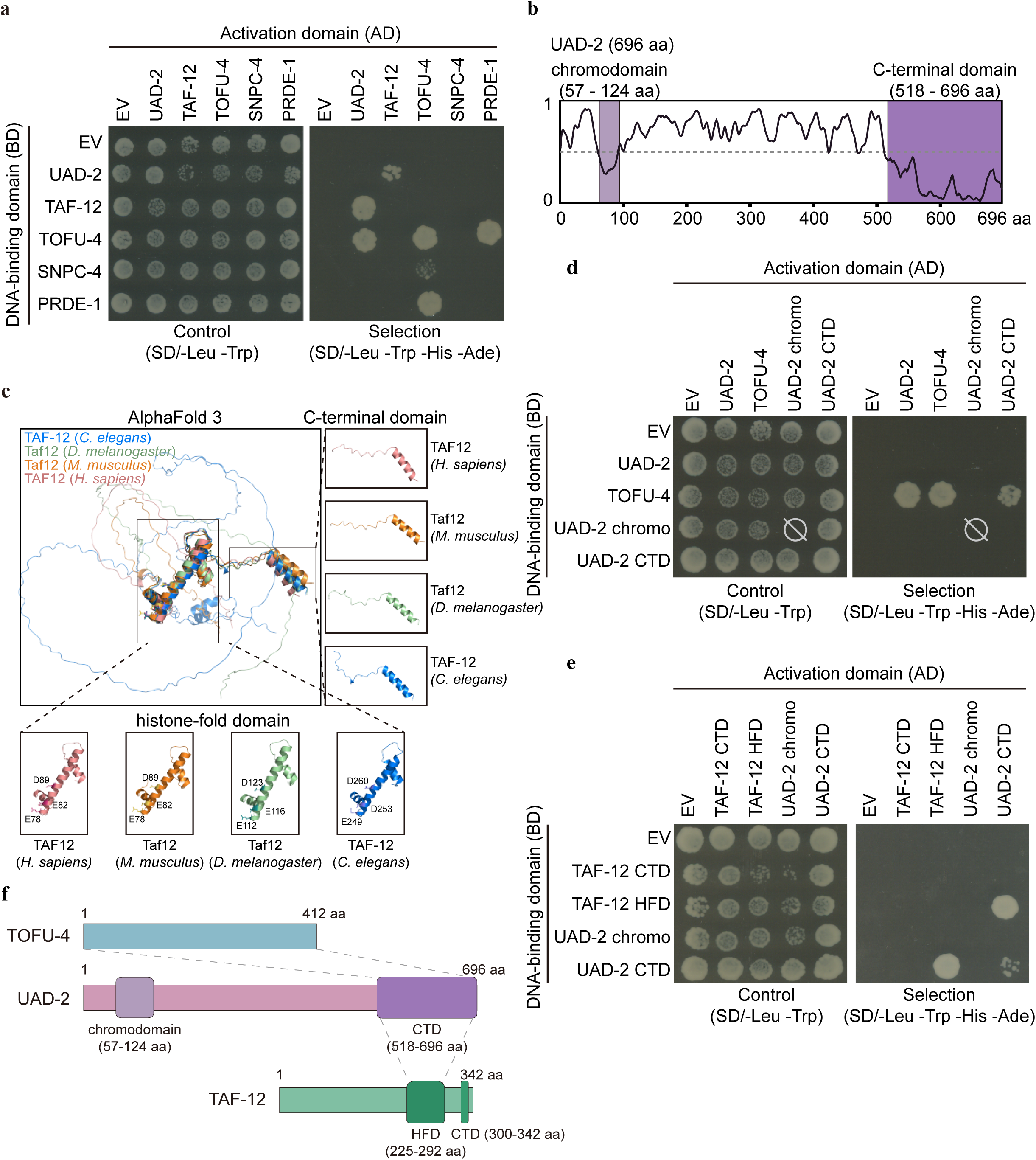
UAD-2 interacts with TOFU-4 and TAF-12 via its C-terminal domain. **a,** Yeast two-hybrid assay to probe protein-protein interactions between UAD-2, TAF-12, and the USTC complex. On nonselective medium (left) all constructs allow growth equally, under selection (right) only strains expressing proteins that interact can grow. Due to the self-activation activity of the DNA binding domain (BD) tethered TOFU-5 protein, TOFU-5 is excluded from the Y2H experiments. **b,** IUPred3 prediction of UAD-2 using the IUPred3 long disorder option. UAD-2 contains a chromodomain and a C-terminal domain. **c,** The AlphaFold 3-predicted structure of TAF-12 of indicated species. TAF-12 encodes a histone-fold domain and a C-terminal domain. **d-e,** Yeast two-hybrid assays are shown probing protein-protein interactions between TOFU-4, the UAD-2 chromodomain (UAD-2 chromo), the UAD-2 C-terminal domain (UAD-2 CTD), the TAF-12 histone-fold domain (TAF-12 HFD), and the TAF-12 C-terminal domain (TAF-12 CTD). TOFU-4 lacks of recognizable domains. **f,** Cartoon of observed protein-protein interactions between TAF-12, UAD-2, and TOFU-4. Data are based on yeast two-hybrid assays using truncation constructs to map interacting domains. CTD: C-terminal domain, HFD: histone-fold domain.

Besides the chromodomain that recognizes H3K27me3, UAD-2 contains a C-terminal domain (CTD) that may facilitate protein-protein interactions, similar to the chromo-shadow domain (Fig. 3b) ^50,63^. TOFU-4 lacks any recognizable domains. Protein sequence alignment revealed that TAF-12 harbors a histone-fold domain (HFD) and a C-terminal domain (CTD), both of which are highly conserved across diverse species (Fig. S3b). The AlphaFold 3-predicted structures of TAF-12 demonstrated that the histone-fold domain and the C-terminal domain of these homologous proteins adopt similar overall structure. The histone-fold domain comprises three tandem α-helices, while the C-terminal domain features a 3_10_ helix (designated as η1), followed by an α-helix (Fig. 3c). The TAF-12 histone-fold domain is crucial for its interaction with the TAF-4 histone-fold domain, which regulates transcriptional activity in germline blastomeres ^64^. Domain mapping experiments showed that the UAD-2 C-terminal domain interacts with both TOFU-4 and the TAF-12 histone-fold domain (Fig. 3d-f and Fig. S3c). Thus, UAD-2 serves as a bridge connecting TAF-12 with the USTC complex.

### The TAF-12 C-terminal domain is essential for piRNA production

TAF-12 contains an intrinsically disordered region (IDR, residues 1-231), a histone-fold domain (HFD, residues 225-292), and a C-terminal domain (CTD, residues 300-342) (Fig. 4a). Using CRISPR-Cas9 technology, we generated *taf-12(ust692/ΔHFD)* and *taf-12(ust550/ΔCTD)* animals (Fig. 4a and Fig. S4a). The homozygous *taf-12(ΔHFD)* animals were embryonically lethal, indicating an essential role of the histone-fold domain. The AlphaFold 3-predicted protein-protein complex structure revealed that the long α4 helix of the TAF-12 histone-fold domain inserts into the angle formed by several α-helices of the UAD-2 C-terminal domain (Fig. 4b, left). The short α3 and α5 helices of the TAF-12 histone-fold domain pack against the α8 helix of the UAD-2 C-terminal domain. Further analysis using PDBsum identified E249, D253, and D260 within the TAF-12 histone-fold domain as key potential interaction residues with the UAD-2 C-terminal domain (Fig. 4b, right). We engineered a *taf-12* mutant with alanine substitutions at these residues (*taf-12(E249A;D253A;D260A)*), which also exhibited embryonic lethality, emphasizing the critical role of these residues for TAF-12 function in addition to engaging in piRNA production.

**Figure 4.**
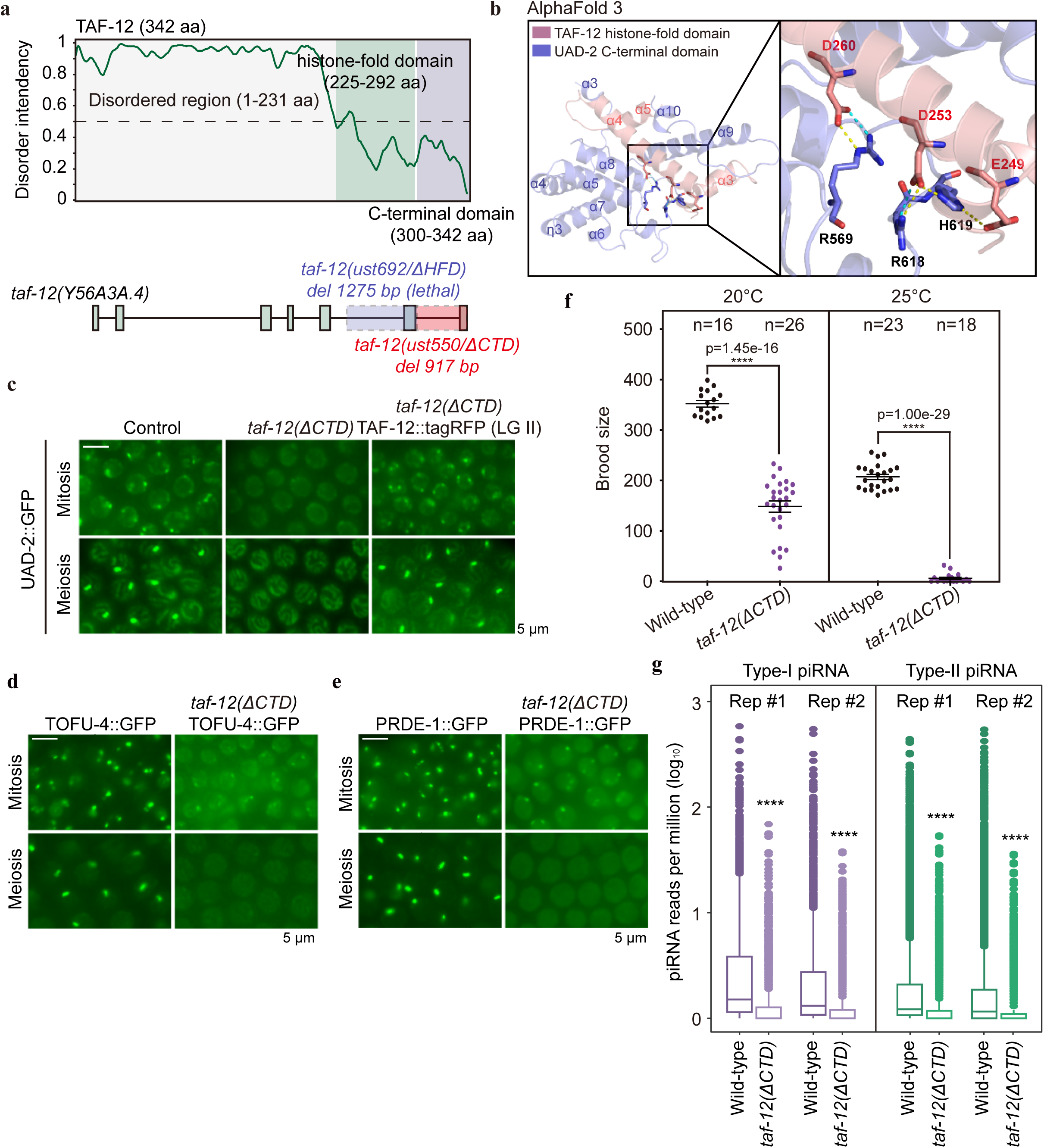
The TAF-12 C-terminal domain is required for piRNA production. **a,** Top: IUPred3 prediction of TAF-12 using the IUPred3 long disorder option. TAF-12 encodes an intrinsically disordered region (1-231 aa). Bottom: schematic of the *taf-12* mutant alleles generated by CRISPR-Cas9 technology. The *taf-12(ust692/ΔHFD)* allele deletes 1275 bp and triggers embryonic lethality. The *taf-12(ust550/ΔCTD)* allele deletes 917 bp and the mutants are viable. **b,** The AlphaFold 3-predicted structure of a TAF-12-UAD-2 complex. The interaction active-site residues of the TAF-12 histone-fold domain and the UAD-2 C-terminal domain are magnified in the bottom right. Salt bridges (yellow) and hydrogen bonds (blue) are depicted with dashed lines. **c,** Fluorescence images showing localization of UAD-2 in mitotic and meiotic germ cells in indicated backgrounds. Scale bar, 5 μm. **d-e,** Mitotic and meiotic germ cells of animals that express TOFU-4::GFP or PRDE-1::GFP in control and *taf-12(ΔCTD)* backgrounds. Scale bar, 5 μm. **f,** Brood size of the indicated animals at 20°C and 25°C, respectively. Bleached embryos were hatched and grown at 20°C or 25°C. Then, L4 stage worms were transferred individually onto fresh NGM plates. A two-tailed *t-test* was performed to determine statistical significance. n > 15. **g,** Boxplots showing type-I and type-II mature piRNA abundance (log_10_ (reads per million)) in wild type and *taf-12(ΔCTD)* mutant worms. Significance was tested with the unpaired Wilcoxon test, ****P < 2.2 x e^−16^. n = 2 biological replicates.

Deletion of the TAF-12 C-terminal domain completely suppressed piRNA focus formation of UAD-2, TOFU-4, and PRDE-1 in meiotic cells and weakened piRNA foci in mitotic cells (Fig. 4c-e). Similar to the TAF-12::AID system, deletion of the TAF-12 C-terminal domain significantly reduced UAD-2::GFP protein levels (Fig. S4b). *taf-12(ΔCTD)* animals displayed reduced brood sizes at both 20°C and 25°C compared to wild-type animals (Fig. 4f), suggesting that the TAF-12 C-terminal domain is essential for fertility. We constructed animals expressing TAF-12(*ΔCTD*)::GFP or TAF-12(*ΔIDR*)::GFP and examined their cellular localization. TAF-12(*ΔIDR*)::GFP still accumulated at piRNA foci, while TAF-12*(ΔCTD)*::GFP did not enrich at piRNA foci throughout the germline (Fig. S4c-e). Thus, the TAF-12 C-terminal domain is crucial for piRNA focus formation and UAD-2 expression.

To test whether the C-terminal domain and the IDR region are essential for TAF-12’s function in piRNA production, we sequenced piRNA populations from wild-type and *taf-12* mutant animals. *taf-12(ΔCTD)* animals showed a significant reduction in both type-I and type-II mature piRNAs (Fig. 4g). Similarly, mature piRNA levels were severely reduced in TAF-12*(ΔCTD)*::GFP mutants (Fig. S4f). In TAF-12(*ΔIDR*)::GFP mutants, piRNA levels were modestly reduced (Fig. S4g).

Taken together, these results suggest that TAF-12 may stabilize the piRNA transcription condensate and promote piRNA transcription through its C-terminal domain.

### TAF-12 collaborates with GTF-2F2, GTF-2H2C, MDT-8, and RPB-5 to promote piRNA transcription

To further investigate the mechanism underlying piRNA transcription, we performed a candidate-based RNAi screen. These candidates included subunits of the Pol II complex, the Mediator complex, the P-TEFb complex, and a set of general transcription factors (TFIIA, -B, -D, -E, -F, and -H) necessary for Pol II-mediated transcription initiation or regulation (Fig. S5a). Among these sixty-five factors, depletion of GTF-2F2, GTF-2H2C, MDT-8, and RPB-5 led to the dispersion of UAD-2 and TAF-12 foci in the nucleus (Fig. 5a-b).

**Figure 5.**
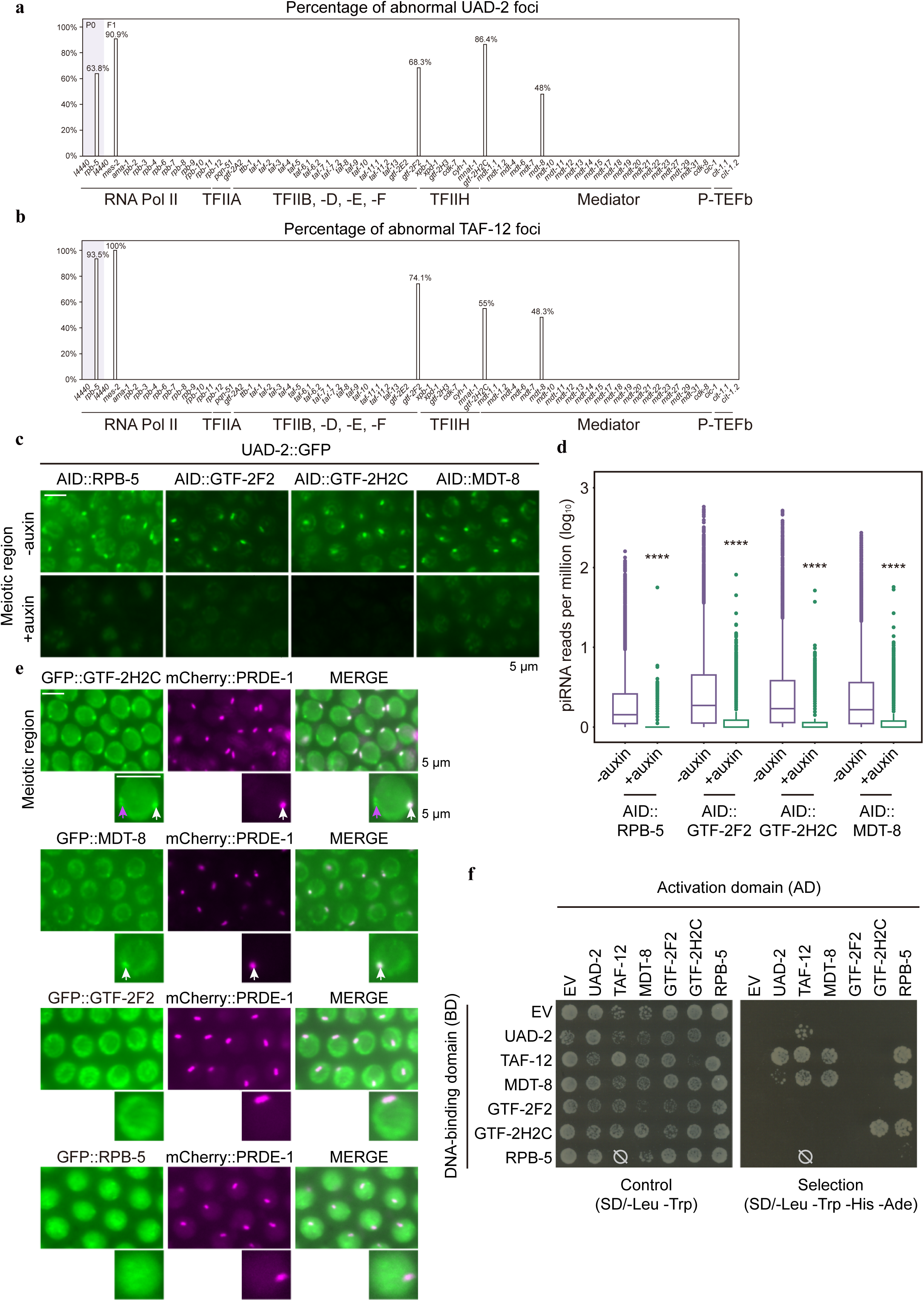
Candidate-based RNAi screen identified that GTF-2F2, GTF-2H2C, MDT-8, and RPB-5 are required for piRNA production. **a-b,** A candidate-based RNAi screen identified transcription factors required for piRNA focus formation of UAD-2 and TAF-12. These candidates include factors of the RNA Pol II, the Mediator complex, the P-TEFb complex and general transcription factors TFIIA, -B, -D, -E, -F, and -H. n > 30. **c,** Fluorescence images of representative meiotic germ cells of the indicated animals expressing UAD-2::GFP and AID::RPB-5, AID::GTF-2F2, AID::GTF-2H2C or AID::MDT-8 (with or without auxin treatment). Scale bar, 5 μm. **d,** The relative abundance of total mature piRNAs in young adult hermaphrodites expressing UAD-2::GFP and AID::RPB-5, AID::GTF-2F2, AID::GTF-2H2C or AID::MDT-8 (with or without auxin treatment). Significance was tested with the unpaired Wilcoxon test, ****P < 2.2 x e^−1^^6^. **e,** Meiotic germ cells of animals that express mCherry::PRDE-1 and GFP::GTF-2H2C, GFP::MDT-8, GFP::GTF-2F2 or GFP::RPB-5. GTF-2H2C and MDT-8 aggregate at piRNA foci and colocalize with PRDE-1. The white arrows indicate piRNA foci, the purple arrow indicates unidentified focus of GFP::GTF-2H2C in germ cells. Scale bar, 5 μm. **f,** Yeast two-hybrid assay to probe for protein-protein interactions between UAD-2, TAF-12, MDT-8, GTF-2F2, GTF-2H2C, and RPB-5. On nonselective medium (left) all constructs allow growth equally, under selection (right) only strains expressing proteins that interact can grow.

GTF-2F2, GTF-2H2C, MDT-8, and RPB-5 are crucial for transcription initiation and PIC assembly ^65–71^. To verify the function of these factors in piRNA focus formation, we generated *in situ* AID-tagged *gtf-2F2*, *gtf-2H2C, mdt-8,* and *rpb-5* animals and confirmed that UAD-2 is unable to form piRNA foci after germline-specific degradation of these factors (Fig. 5c).

We sequenced small RNA populations from control and auxin-treated animals. Germline-specific degradation of GTF-2F2, GTF-2H2C, MDT-8, and RPB-5 resulted in a significant reduction of mature piRNAs (Fig. 5d and Fig. S5b-e). We then generated animals expressing GFP-tagged GTF-2F2, GTF-2H2C, MDT-8, or RPB-5 and examined their cellular localization. These proteins were broadly expressed in somatic nuclei (Fig. S5f). In germ cells, GTF-2F2 and RPB-5 did not exhibit strong focus formation, whereas MDT-8 and GTF-2H2C accumulated in piRNA foci and colocalized with PRDE-1 (Fig. 5e). Notably, GTF-2H2C formed an additional, unidentified focus in germ cells, which was distinct from the piRNA foci (Fig. 5e).

Using yeast two-hybrid assays, we examined protein-protein interactions among UAD-2, TAF-12, GTF-2F2, GTF-2H2C, MDT-8, and RPB-5. The analyses revealed that TAF-12 interacts with MDT-8 and RPB-5 (Fig. 5f and Fig. S5g). MDT-8 and RPB-5 also interact with each other. Interestingly, MDT-8 showed a moderate interaction with UAD-2 (Fig. 5f and Fig. S5g). Thus, we concluded that TAF-12 bridges UAD-2 and the transcription machinery to promote piRNA transcription.

## Discussion

In this study, we demonstrated that TAF-12 serves as a crucial molecular link between the USTC complex, UAD-2, and the general transcription machinery at piRNA clusters, which are typically enriched with the facultative histone mark H3K27me3 ^26,33^. TAF-12 forms piRNA foci in germ cells, and this aggregation relies on both UAD-2 and the USTC complex. The knockdown of TAF-12 results in the loss of both piRNA precursors and mature piRNAs. TAF-12 may initiate piRNA transcription by recruiting GTF-2F2, GTF-2H2C, MDT-8, and RPB-5. Our findings indicate a large piRNA transcription complex comprised of TAF-12, UAD-2, the USTC complex, three general transcription factors, and a subunit of the RNA Pol II (Fig. 6). The USTC complex associates with the Ruby motif located upstream of piRNA transcription start sites (TSSs). TOFU-4, a component of the USTC complex, recruits UAD-2 to piRNA genes. UAD-2 recognizes H3K27me3 marks across piRNA clusters via its chromodomain and interacts with TAF-12 through its C-terminal domain. This interaction facilitates the subsequent recruitment of the piRNA transcription machinery.

**Figure 6.**
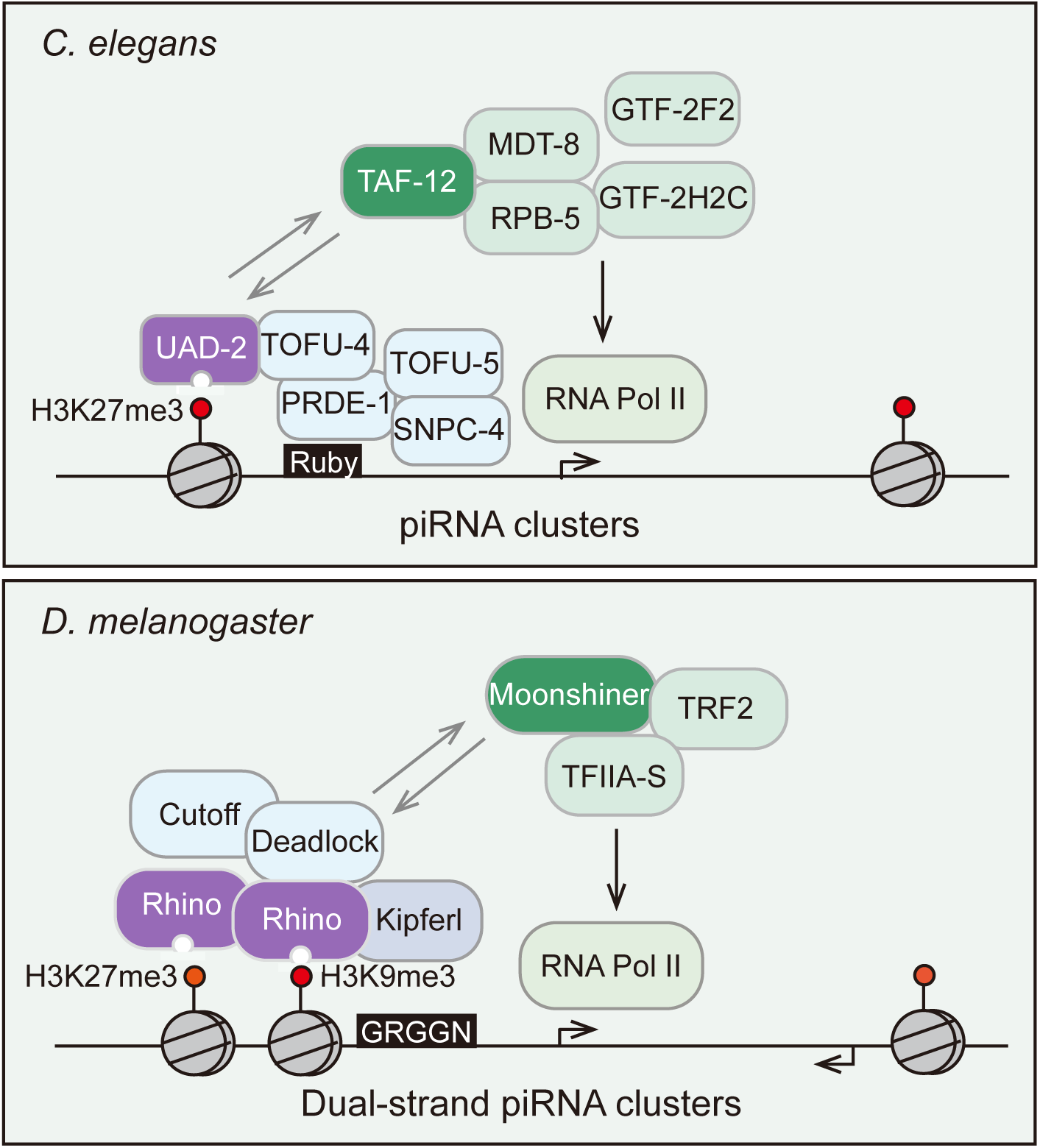
Schematic model illustrating piRNA transcription from heterochromatic regions. (Top) In *C. elegans*, the transcription of piRNA genes from piRNA clusters is facilitated by the recognition of the heterochromatic histone mark H3K27me3 via the chromodomain protein UAD-2. UAD-2 interacts with the upstream sequence transcription complex (USTC) and recruits the TFIID component TAF-12 to initiate the transcription of piRNA precursors. (Bottom) Similarly, in *D. melanogaster*, piRNA transcription from dual-strand piRNA clusters relies on the reader protein of the heterochromatic histone mark H3K9me3 and H3K27me3, Rhino. Deadlock interacts with Rhino and recruits Moonshiner, TRF2, and TFIIA-S to facilitate piRNA transcription. Kipferl binds to G-rich DNA motifs and interacts with Rhino, thereby influencing Rhino’s binding profile.

Our data reveal that piRNA transcription in heterochromatic regions relies on histone modification readers to recruit a specific transcription machinery. Similarly, in *S. pombe*, the transcription of siRNAs from heterochromatic loci depends on the chromodomain protein Chp1 ^72–74^. Chp1 binds to methylated H3K9 via its chromodomain and tethers the RNAi-induced transcriptional silencing complex (RITS) to heterochromatin, thereby promoting siRNA production ^75–77^. In *Arabidopsis thaliana,* the initial transcription of 24-nt siRNAs is mediated by SHH1 ^78,79^. SHH1 binds to H3K9me2 via its SAWADEE domain and recruits Pol IV to transcribe precursor RNAs ^80–82^. In *Drosophila* ovarian germ cells, the chromodomain protein Rhino specifically binds to piRNA source loci enriched with H3K9me3 and collaborates with Deadlock, Cutoff, and a Moonshiner-dependent transcription machinery to initiate non-canonical transcription ^83^. These findings suggest that eukaryotes may employ a universally conserved strategy to facilitate small RNA production in heterochromatic regions.

Recent studies in *Drosophila* have shown that H3K9me3 and H3K27me3 modifications coexist at several Rhino-dependent loci ^55^. At these loci, both H3K9me3 and H3K27me3 are necessary for piRNA transcription. In contrast to *yeast*, *plants* and *Drosophila*, H3K27me3 is essential for piRNA production in *C. elegans*, while H3K9 methylation appears to be dispensable for piRNA biogenesis. The *met-2;set-25* double mutation, which completely disrupts mono-, di-, and trimethylation of H3K9, shows no observable impact on piRNA focus formation ^30,84,85^. It will be interesting to investigate why the H3K27me3 mark is critical for piRNA transcription and how H3K27me3 marks are established at piRNA clusters.

There has been a long-standing debate over how the USTC complex is initially recruited to piRNA clusters. Early models proposed that the USTC complex binds to the Ruby motif upstream of type I piRNA genes, yet currently lacking evidence that TOFU-4, TOFU-5, PRDE-1, or SNPC-4 directly binds to chromatin. In *Drosophila*, Kipferl encodes two zinc finger arrays and specifically binds to guanosine-rich DNA motifs ^49^. Kipferl interacts with the Rhino chromodomain to define Rhino’s binding profile within H3K9me2- and H3K9me3-enriched domains. It will be interesting to investigate whether ZnF proteins in *C. elegans* employ a similar specialized mechanism to recruit the USTC complex at piRNA genes. Additionally, TOFU-4 and TAF-12 interact with the UAD-2 C-terminal domain, suggesting that TOFU-4 may compete with TAF-12 for binding to UAD-2 proteins. Detailed structural analysis of piRNA genes, the USTC complex, UAD-2, and TAF-12 could provide critical insights into the mechanism of piRNA transcription.

## Materials and Methods

### Strains

The Bristol strain N2 was used as the standard wild-type strain. All strains were grown at 20°C unless otherwise specified. The strains used in this study are listed in Table S1.

### Construction of plasmids and transgenic strains

For the transgenes of transcription factors, endogenous promoter sequences, UTRs, and ORFs of transcription factors were PCR-amplified with the primers listed in Table S2. A *gfp::3xflag* fused to a linker sequence (GGAGGTGGAGGTGGAGCT) was PCR-amplified with the primers 5′-GGAGGTGGAGGTGGAGCTAT −3′ and 5′-CTTGTCATCGTCATCCTTGTAATCGA −3′ from SHG1093 genomic DNA. A *tagRFP::3xha* fused to a linker sequence (GGAGGTGGAGGTGGAGCT) was PCR-amplified with the primers 5′-GGAGGTGGAGGTGGAGCTATG −3′ and 5′-GTAATCTGGAACATCGTATGGGTAAGCGTAATCTGGAACATCGTATGGGTAGTTGAGCTTGTGCCCG −3′ from SHG2673 (*rrf-1(ust364[rrf-1::tagRFP]) I*) genomic DNA ^40^. ClonExpress MultiS One Step Cloning Kit (Vazyme C113-02, Nanjing) was used to connect these fragments with vector which is amplified with 5′-TGTGAAATTGTTATCCGCTGGT −3′ and 5′-CACACGTGCTGGCGTTAC −3′ from pCFJ151. The injection mix contained PDD162 (50 ng/µl), transcription factors repair plasmid (50 ng/µl), pCFJ90 (5 ng/µl), and three sgRNAs (30 ng/µl). The mix was injected into young adult N2 animals. The transcription factors’ transgenes were integrated into the *C. elegans* genome locus of each gene in situ via a multiple sgRNA-based CRISPR-Cas9 gene editing system ^86^.

### Construction of deletion mutants

For gene deletions, triple-sgRNA-guided chromosome deletion was conducted as previously described ^86^. To construct sgRNA expression vectors, the 20 bp *unc-119* sgRNA guide sequence in the *pU6::unc-119* sgRNA(F+E) vector was replaced with different sgRNA guide sequences. Addgene plasmid #47549 was utilized to express the Cas9 II protein. Plasmid mixtures containing 30 ng/μl of each of the three or four sgRNA expression vectors, 50 ng/μl of the Cas9 II expressing plasmid, and 5 ng/μl pCFJ90 were co-injected into N2 animals. Deletion mutants were screened by PCR amplification and confirmed by sequencing. The sgRNA sequences are listed in Table S3.

### Immunoprecipitation followed by mass spectrometry analysis

IP-MS was conducted as previously reported ^36^. Mixed-stage transgenic worms expressing UAD-2::GFP were collected and resuspended in equal volumes of 2x lysis buffer (50 mM Tris-HCl [pH 8.0], 300 mM NaCl, 10% glycerol, 1% Triton X-100, Roche ®cOmplete EDTA-Free Protease Inhibitor Cocktail, 10 mM NaF, and 2 mM Na_3_VO_4_) and lysed using a FastPrep-24 5G homogenizer. The lysate supernatant was incubated with in-house-prepared anti-GFP beads for one hour at 4°C. The beads were then washed three times with cold lysis buffer. The GFP immunoprecipitates were eluted with chilled elution buffer (100 mM glycine-HCl [pH 2.5]). Approximately 1/8 of each eluate was subjected to western blot analysis, while the reminder was precipitated with TCA or cold acetone and dissolved in 100 mM Tris (pH 8.5) containing 8 M urea. Proteins were reduced with TCEP, alkylated with IAA, and finally digested with trypsin at 37°C overnight. LC–MS/MS analysis of the resulting peptides and MS data processing were performed as previously described ^87^. A WD scoring matrix was employed to identify high-confidence candidate interacting proteins ^87^.

### Western blotting

Synchronized young adult worms incubated at 20°C were collected and washed three times with 1x M9 buffer. Samples were stored at −80°C until use. The worms were suspended in 1x SDS loading buffer and heated in a metal bath at 95°C for 5 - 10 minutes. The suspensions were then centrifuged at 17,000 g, and the supernatants were collected. Proteins were resolved by SDS-PAGE on gradient gels (10% separation gel, 5% spacer gel) and transferred to a Hybond-ECL membrane. After washing with 1x Tris-buffered saline with Tween-20 (TBST) buffer and blocking with 5% milk-TBST, the membrane was incubated overnight at 4°C with antibodies. The membrane was washed three times for 10 minutes each with 1x TBST and then incubated with secondary antibodies at room temperature for 2 hours. The membrane was washed thrice for 10 min with 1x TBST and then visualized. Antibodies used for western blotting: anti-FLAG (Sigma, F1804), 1:1000; anti-β-actin (Beyotime, AF5003), 1:5000. Secondary antibodies: HRP-labeled goat anti-rabbit IgG (H+L) (Abcam ab205718), 1:15000; HRP-labeled goat anti-mouse IgG (H+L) (Beyotime A0216), 1:1000.

### RNA isolation

Young adult synchronized populations of worms were grown and washed three times with 1x M9 to remove bacterial residues. For RNA extraction, 400 µl of TRIzol Reagent (Ambion, 15596026) was added to a 100 µl worm aliquot, followed by 7-8 cycles of freezing in liquid nitrogen and thawing in a 42°C water bath. Afterward, 100 µl of DNA/RNA Extraction Reagent (Solarbio LIFE SCIENCES, P1014) was added to the samples, which were then centrifuged for 15 minutes at 13,200 rpm at 4 °C. The supernatant was further treated with 400 µl isopropanol, 400 µl pre-chilled 75% ethanol, and subjected to DNase I digestion (Thermo Fisher Scientific, EN0521). RNA was eluted into 20 µl of nuclease-free water, and each sample was divided into three aliquots for piRNA precursor, mature piRNA, and mRNA library preparation.

### FastAP/RppH treatment for piRNA precursors

FastAP treatment of 2 µg of isolated RNA was performed in FastAP buffer using 2 µl of FastAP (Thermo Fisher Scientific^TM^, EP0654) in a 40 µl reaction. The reaction was incubated at 37°C for 10 minutes, followed by heat inactivation for 5 minutes at 75°C. The FastAP-treated RNA was extracted and precipitated overnight, following the protocol for decapping treatment with RppH (NEB, #M0356) according to the vendor’s instructions. The resulting RNA was used as input for small RNA library preparation.

### Library preparation and sequencing

Small RNAs were subjected to RNA deep sequencing using an Illumina platform (AZENTA LIFE SCIENCES). Briefly, small RNAs ranging from 15 to 40 nucleotides were gel-purified and ligated to a P7 adapter (5′-AGATCGGAAGAGCACACGTCTGAACTCCAGTCAC −3′) and a P5 adapter (5′-AGATCGGAAGAGCGTCGTGTAGGGAAAGAGTGT −3′). The ligation products were gel-purified, reverse transcribed, amplified, and sequenced using an Illumina Novaseq platform.

### Small RNA-seq analysis

The Illumina-generated raw reads were first filtered to remove adapters, low-quality tags, and contaminants to obtain clean reads using fastp. For mature piRNA and pre-piRNA analysis, clean reads ranging from 17 to 35 nt were respectively mapped to mature piRNA regions, pre-piRNA regions, and the *C. elegans* transcriptome assembly WS243 using Bowtie2 with default parameters. The number of reads targeting each transcript was counted using custom Perl scripts. The number of total reads mapped to the transcriptome minus the number of total reads corresponding to sense ribosomal RNA (rRNA) transcripts (5S, 5.8S, 18S, and 26S), and sense protein coding mRNA reads were used as the normalization number to exclude the possible degradation fragments of sense rRNAs and mRNAs.

### piRNA gene annotations

piRNA annotations were downloaded from the piRBase online database (http://www.regulatoryrna.org/database/piRNA). Genomic coordinates of piRNA genes were obtained by SAMtools against the *C. elegans* ce11 genome assembly. Type-II piRNA genes were obtained from a previous publication ^19^. Type-I piRNA gene lists were created by filtering the piRBase annotations with type-II piRNA genes.

### mRNA-seq analysis

The Illumina-generated raw reads were first filtered to remove adapters, low-quality tags, and contaminants to obtain clean reads at Novogene. The clean reads were mapped to the reference genome of WBcel235 via HISAT2 software (version 2.1.0) ^88^. Differential expression analysis was performed using custom R scripts. A twofold-change cutoff was applied when filtering for differentially expressed genes. All plots were drawn using custom R scripts.

### Candidate-based RNAi screen

RNAi experiments were performed at 20°C by placing synchronized embryos on feeding plates as previously described ^30^. HT115 bacteria expressing the empty vector L4440 were used as negative controls. Bacterial clones expressing double-stranded RNAs (dsRNAs) were obtained from the Ahringer RNAi library and were sequenced to verify their identity. All feeding RNAi experiments were performed for two generations except for sterile worms, which were RNAi treated for one generation. Images were collected using a Leica DM4 B microscope.

### Yeast Two-Hybrid assay

The yeast two-hybrid assay was performed according to the manufacturer’s protocol (Clontech user manual 630489). Briefly, the *Saccharomyces cerevisiae* strain AH109 was grown in YPDA selective medium at 30℃. Assayed proteins were fused to the activation domain (AD) and DNA-binding domain (BD) of the Gal4 transcription factor and transformed into pGADT7 (AD) and pGBKT7 (BD). Individual colonies of transformed haploids were selected, picked and mated. Upon interaction of two fusion proteins the Gal4 transcription factor is reconstituted and will activate the reporter genes (ADE2, HIS3), which will allow growth on synthetic defined medium lacking Ade, His, Leu and Trp.

### Brood Size

L4 hermaphrodites were singled onto plates and transferred daily as adults until embryo production ceased and the progeny numbers were scored.

### Statistics

Bar graphs with error bars are presented with mean and standard deviation (SD). All experiments were conducted with independent *C. elegans* animals for the indicated N times. Statistical analysis was performed with two-tailed Student’s t test.

## Acknowledgments

We are grateful to the members of the Guang laboratory for their comments and to Dr. Yong Ding’s laboratory for yeast strains. We are grateful to the International *C. elegans* Gene Knockout Consortium and the National Bioresource Project for providing the strains. Some strains were provided by the CGC, which is funded by the NIH Office of Research Infrastructure Programs (P40 OD010440).

## Author contributions

S.G., C.Z. and X.H. conceptualized the research; X.F., X.C., Y.Z., C.Z., and S.G. designed the research; X.H., Y.K., M.L., M.H., J.C., W.W., T.H., and X.S. performed the research; X.H., Y.K., M.L., J.C., T.H., and X.S. contributed new reagents; X.H., Y.K., and M.H. provided analytic tools and performed bioinformatics analysis; X.H., X.C., X.F., and S.G. wrote the paper.

## Data Availability

The raw sequence data reported in this paper have been deposited in the Genome Sequence Archive in the National Genomics Data Center, China National Center for Bioinformation / Beijing Institute of Genomics, Chinese Academy of Sciences (GSA: CRA022192) and are publicly accessible at https://ngdc.cncb.ac.cn/gsa/s/u347JF83.

## References

1. Mao, H., Zhu, C., Zong, D., Weng, C., Yang, X., Huang, H., Liu, D., Feng, X., and Guang, S. (2015). The Nrde Pathway Mediates Small-RNA-Directed Histone H3 Lysine 27 Trimethylation in Caenorhabditis elegans. Curr Biol 25, 2398–2403. 10.1016/j.cub.2015.07.051.

2. Batista, P.J., Ruby, J.G., Claycomb, J.M., Chiang, R., Fahlgren, N., Kasschau, K.D., Chaves, D.A., Gu, W., Vasale, J.J., Duan, S., et al. (2008). PRG-1 and 21U-RNAs interact to form the piRNA complex required for fertility in C. elegans. Mol Cell 31, 67–78. 10.1016/j.molcel.2008.06.002.

3. Lee, H.C., Gu, W.F., Shirayama, M., Youngman, E., Conte, D., and Mello, C.C. (2012). C. elegans piRNAs Mediate the Genome-wide Surveillance of Germline Transcripts. Cell 150, 78–87. 10.1016/j.cell.2012.06.016.

4. Shen, E.Z., Chen, H., Ozturk, A.R., Tu, S.K., Shirayama, M., Tang, W., Ding, Y.H., Dai, S.Y., Weng, Z.P., and Mello, C.C. (2018). Identification of piRNA Binding Sites Reveals the Argonaute Regulatory Landscape of the Germline. Cell 172, 937–951. 10.1016/j.cell.2018.02.002.

5. Vagin, V.V., Sigova, A., Li, C., Seitz, H., Gvozdev, V., and Zamore, P.D. (2006). A distinct small RNA pathway silences selfish genetic elements in the germline. Science 313, 320–324. 10.1126/science.1129333.

6. Aravin, A., Gaidatzis, D., Pfeffer, S., Lagos-Quintana, M., Landgraf, P., Iovino, N., Morris, P., Brownstein, M.J., Kuramochi-Miyagawa, S., Nakano, T., et al. (2006). A novel class of small RNAs bind to MILI protein in mouse testes. Nature 442, 203–207. 10.1038/nature04916.

7. Girard, A., Sachidanandam, R., Hannon, G.J., and Carmell, M.A. (2006). A germline-specific class of small RNAs binds mammalian Piwi proteins. Nature 442, 199–202. 10.1038/nature04917.

8. Kiuchi, T., Koga, H., Kawamoto, M., Shoji, K., Sakai, H., Arai, Y., Ishihara, G., Kawaoka, S., Sugano, S., Shimada, T., et al. (2014). A single female-specific piRNA is the primary determiner of sex in the silkworm. Nature 509, 633–636. 10.1038/nature13315.

9. Schnettler, E., Donald, C.L., Human, S., Watson, M., Siu, R.W.C., McFarlane, M., Fazakerley, J.K., Kohl, A., and Fragkoudis, R. (2013). Knockdown of piRNA pathway proteins results in enhanced Semliki Forest virus production in mosquito cells. J Gen Virol 94, 1680–1689. 10.1099/vir.0.053850-0.

10. Grentzinger, T., Armenise, C., Brun, C., Mugat, B., Serrano, V., Pelisson, A., and Chambeyron, S. (2012). piRNA-mediated transgenerational inheritance of an acquired trait. Genome Res 22, 1877–1888. 10.1101/gr.136614.111.

11. Shirayama, M., Seth, M., Lee, H.C., Gu, W., Ishidate, T., Conte, D., Jr., and Mello, C.C. (2012). piRNAs initiate an epigenetic memory of nonself RNA in the C. elegans germline. Cell 150, 65–77. 10.1016/j.cell.2012.06.015.

12. Ashe, A., Sapetschnig, A., Weick, E.M., Mitchell, J., Bagijn, M.P., Cording, A.C., Doebley, A.L., Goldstein, L.D., Lehrbach, N.J., Le Pen, J., et al. (2012). piRNAs Can Trigger a Multigenerational Epigenetic Memory in the Germline of C. elegans. Cell 150, 88–99. 10.1016/j.cell.2012.06.018.

13. Luteijn, M.J., van Bergeijk, P., Kaaij, L.J., Almeida, M.V., Roovers, E.F., Berezikov, E., and Ketting, R.F. (2012). Extremely stable Piwi-induced gene silencing in Caenorhabditis elegans. EMBO J 31, 3422–3430. 10.1038/emboj.2012.213.

14. Brennecke, J., Malone, C.D., Aravin, A.A., Sachidanandam, R., Stark, A., and Hannon, G.J. (2008). An epigenetic role for maternally inherited piRNAs in transposon silencing. Science 322, 1387–1392. 10.1126/science.1165171.

15. Wang, G., and Reinke, V. (2008). A C. elegans Piwi, PRG-1, regulates 21U-RNAs during spermatogenesis. Curr Biol 18, 861-867. 10.1016/j.cub.2008.05.009.

16. Yigit, E., Batista, P.J., Bei, Y.X., Pang, K.M., Chen, C.C.G., Tolia, N.H., Joshua-Tor, L., Mitani, S., Simard, M.J., and Mello, C.C. (2006). Analysis of the argonaute family reveals that distinct argonautes act sequentially during RNAi. Cell 127, 747–757. 10.1016/j.cell.2006.09.033.

17. Das, P.P., Bagijn, M.P., Goldstein, L.D., Woolford, J.R., Lehrbach, N.J., Sapetschnig, A., Buhecha, H.R., Gilchrist, M.J., Howe, K.L., Stark, R., et al. (2008). Piwi and piRNAs act upstream of an endogenous siRNA pathway to suppress Tc3 transposon mobility in the Caenorhabditis elegans germline. Mol Cell 31, 79–90. 10.1016/j.molcel.2008.06.003.

18. Bagijn, M.P., Goldstein, L.D., Sapetschnig, A., Weick, E.M., Bouasker, S., Lehrbach, N.J., Simard, M.J., and Miska, E.A. (2012). Function, targets, and evolution of Caenorhabditis elegans piRNAs. Science 337, 574–578. 10.1126/science.1220952.

19. Gu, W., Lee, H.C., Chaves, D., Youngman, E.M., Pazour, G.J., Conte, D., Jr., and Mello, C.C. (2012). CapSeq and CIP-TAP identify Pol II start sites and reveal capped small RNAs as C. elegans piRNA precursors. Cell 151, 1488–1500. 10.1016/j.cell.2012.11.023.

20. Ruby, J.G., Jan, C., Player, C., Axtell, M.J., Lee, W., Nusbaum, C., Ge, H., and Bartel, D.P. (2006). Large-scale sequencing reveals 21U-RNAs and additional microRNAs and endogenous siRNAs in C. elegans. Cell 127, 1193–1207. 10.1016/j.cell.2006.10.040.

21. Billi, A.C., Freeberg, M.A., Day, A.M., Chun, S.Y., Khivansara, V., and Kim, J.K. (2013). A Conserved Upstream Motif Orchestrates Autonomous, Germline-Enriched Expression of piRNAs. Plos Genet 9, e1003392. 10.1371/journal.pgen.1003392.

22. Weng, C., Kosalka, J., Berkyurek, A.C., Stempor, P., Feng, X., Mao, H., Zeng, C., Li, W.J., Yan, Y.H., Dong, M.Q., et al. (2019). The USTC co-opts an ancient machinery to drive piRNA transcription in C. elegans. Gene Dev 33, 90–102. 10.1101/gad.319293.118.

23. Weick, E.M., Sarkies, P., Silva, N., Chen, R.A., Moss, S.M.M., Cording, A.C., Ahringer, J., Martinez-Perez, E., and Miska, E.A. (2014). PRDE-1 is a nuclear factor essential for the biogenesis of Ruby motif-dependent piRNAs in C. elegans. Gene Dev 28, 783–796. 10.1101/gad.238105.114.

24. Kasper, D.M., Wang, G.L., Gardner, K.E., Johnstone, T.G., and Reinke, V. (2014). The SNAPc Component SNPC-4 Coats piRNA Domains and Is Globally Required for piRNA Abundance. Dev Cell 31, 145–158. 10.1016/j.devcel.2014.09.015.

25. Zhang, G., Zheng, C., Ding, Y.H., and Mello, C.C. (2024). Casein kinase II promotes piRNA production through direct phosphorylation of USTC component TOFU-4. Nat Commun 15, 2727. 10.1038/s41467-024-46882-9.

26. Beltran, T., Barroso, C., Birkle, T.Y., Stevens, L., Schwartz, H.T., Sternberg, P.W., Fradin, H., Gunsalus, K., Piano, F., Sharma, G., et al. (2019). Comparative Epigenomics Reveals that RNA Polymerase II Pausing and Chromatin Domain Organization Control Nematode piRNA Biogenesis. Dev Cell 48, 793–810. 10.1016/j.devcel.2018.12.026.

27. Gaydos, L.J., Wang, W.C., and Strome, S. (2014). H3K27me and PRC2 transmit a memory of repression across generations and during development. Science 345, 1515–1518. 10.1126/science.1255023.

28. Ketel, C.S., Andersen, E.F., Vargas, M.L., Suh, J., Strome, S., and Simon, J.A. (2005). Subunit contributions to histone methyltransferase activities of fly and worm polycomb group complexes. Mol Cell Biol 25, 6857–6868. 10.1128/MCB.25.16.6857-6868.2005.

29. Yuzyuk, T., Fakhouri, T.H.I., Kiefer, J., and Mango, S.E. (2009). The Polycomb Complex Protein Promotes the Transition from Developmental Plasticity to Differentiation in Embryos. Dev Cell 16, 699–710. 10.1016/j.devcel.2009.03.008.

30. Huang, X., Cheng, P., Weng, C., Xu, Z., Zeng, C., Xu, Z., Chen, X., Zhu, C., Guang, S., and Feng, X. (2021). A chromodomain protein mediates heterochromatin-directed piRNA expression. Proc Natl Acad Sci U S A 118, e2103723118. 10.1073/pnas.2103723118.

31. Hou, X., Xu, M., Zhu, C., Gao, J., Li, M., Chen, X., Sun, C., Nashan, B., Zang, J., Zhou, Y., et al. (2023). Systematic characterization of chromodomain proteins reveals an H3K9me1/2 reader regulating aging in C. elegans. Nature Communications 14, 1254. 10.1038/s41467-023-36898-y.

32. Zhu, C., Si, X., Hou, X., Xu, P., Gao, J., Tang, Y., Weng, C., Xu, M., Yan, Q., Jin, Q., et al. (2023). Spatially clustered piRNA genes promote the transcription of piRNAs via condensate formation of the H3K27me3 reader UAD-2. bioRxiv 2023, 12.10.571043. 10.1101/2023.12.10.571043.

33. Paniagua, N., Roberts, C.J., Gonzalez, L.E., Monedero-Alonso, D., and Reinke, V. (2024). The Upstream Sequence Transcription Complex dictates nucleosome positioning and promoter accessibility at piRNA genes in the germ line. Plos Genet 20, e1011345. 10.1371/journal.pgen.1011345.

34. Berkyurek, A.C., Furlan, G., Lampersberger, L., Beltran, T., Weick, E.M., Nischwitz, E., Cunha Navarro, I., Braukmann, F., Akay, A., Price, J., et al. (2021). The RNA polymerase II subunit RPB-9 recruits the integrator complex to terminate Caenorhabditis elegans piRNA transcription. EMBO J 40, e105565. 10.15252/embj.2020105565.

35. Beltran, T., Pahita, E., Ghosh, S., Lenhard, B., and Sarkies, P. (2021). Integrator is recruited to promoter-proximally paused RNA Pol II to generate piRNA precursors. EMBO J 40, e105564. 10.15252/embj.2020105564.

36. Zeng, C., Weng, C., Wang, X., Yan, Y.H., Li, W.J., Xu, D., Hong, M., Liao, S., Dong, M.Q., Feng, X., et al. (2019). Functional Proteomics Identifies a PICS Complex Required for piRNA Maturation and Chromosome Segregation. Cell Rep 27, 3561–3572. 10.1016/j.celrep.2019.05.076.

37. Rodrigues, R.J.C., Domingues, A.M.D., Hellmann, S., Dietz, S., de Albuquerque, B.F.M., Renz, C., Ulrich, H.D., Sarkies, P., Butter, F., and Ketting, R.F. (2019). PETISCO is a novel protein complex required for 21U RNA biogenesis and embryonic viability. Gene Dev 33, 857–870. 10.1101/gad.322446.118.

38. Wang, X., Zeng, C., Liao, S., Zhu, Z., Zhang, J., Tu, X., Yao, X., Feng, X., Guang, S., and Xu, C. (2021). Molecular basis for PICS-mediated piRNA biogenesis and cell division. Nat Commun 12, 5595. 10.1038/s41467-021-25896-7.

39. Perez-Borrajero, C., Podvalnaya, N., Holleis, K., Lichtenberger, R., Karaulanov, E., Simon, B., Basquin, J., Hennig, J., Ketting, R.F., and Falk, S. (2021). Structural basis of PETISCO complex assembly during piRNA biogenesis in C. elegans. Genes Dev 35, 1304–1323. 10.1101/gad.348648.121.

40. Huang, X., Feng, X., Yan, Y.H., Xu, D., Wang, K., Zhu, C., Dong, M.Q., Huang, X., Guang, S., and Chen, X. (2024). Compartmentalized localization of perinuclear proteins within germ granules in C. elegans. Dev Cell 60, 1–20. 10.1016/j.devcel.2024.12.016.

41. Podvalnaya, N., Bronkhorst, A.W., Lichtenberger, R., Hellmann, S., Nischwitz, E., Falk, T., Karaulanov, E., Butter, F., Falk, S., and Ketting, R.F. (2023). piRNA processing by a trimeric Schlafen-domain nuclease. Nature 622, 402–409. 10.1038/s41586-023-06588-2.

42. Tang, W., Tu, S., Lee, H.C., Weng, Z.P., and Mello, C.C. (2016). The RNase PARN-1 Trims piRNA 3’ Ends to Promote Transcriptome Surveillance in C. elegans. Cell 164, 974–984. 10.1016/j.cell.2016.02.008.

43. Montgomery, T.A., Rim, Y.S., Zhang, C., Dowen, R.H., Phillips, C.M., Fischer, S.E.J., and Ruvkun, G. (2012). PIWI Associated siRNAs and piRNAs Specifically Require the Caenorhabditis elegans HEN1 Ortholog henn-1. Plos Genet 8, e1002616. 10.1371/journal.pgen.1002616.

44. Billi, A.C., Alessi, A.F., Khivansara, V., Han, T., Freeberg, M., Mitani, S., and Kim, J.K. (2012). The Caenorhabditis elegans HEN1 ortholog, HENN-1, methylates and stabilizes select subclasses of germline small RNAs. Plos Genet 8, e1002617. 10.1371/journal.pgen.1002617.

45. Kamminga, L.M., van Wolfswinkel, J.C., Luteijn, M.J., Kaaij, L.J.T., Bagijn, M.P., Sapetschnig, A., Miska, E.A., Berezikov, E., and Ketting, R.F. (2012). Differential Impact of the HEN1 Homolog HENN-1 on 21U and 26G RNAs in the Germline of Caenorhabditis elegans. Plos Genet 8, e1002702. 10.1371/journal.pgen.1002702.

46. Brennecke, J., Aravin, A.A., Stark, A., Dus, M., Kellis, M., Sachidanandam, R., and Hannon, G.J. (2007). Discrete small RNA-generating loci as master regulators of transposon activity in Drosophila. Cell 128, 1089–1103. 10.1016/j.cell.2007.01.043.

47. Klattenhoff, C., Xi, H.L., Li, C.J., Lee, S., Xu, J., Khurana, J.S., Zhang, F., Schultz, N., Koppetsch, B.S., Nowosielska, A., et al. (2009). The HP1 Homolog Rhino Is Required for Transposon Silencing and piRNA Production by Dual-Strand Clusters. Cell 138, 1137–1149. 10.1016/j.cell.2009.07.014.

48. Vermaak, D., and Malik, H.S. (2009). Multiple roles for heterochromatin protein 1 genes in Drosophila. Annu Rev Genet 43, 467–492. 10.1146/annurev-genet-102108-134802.

49. Baumgartner, L., Handler, D., Platzer, S.W., Yu, C., Duchek, P., and Brennecke, J. (2022). The Drosophila ZAD zinc finger protein Kipferl guides Rhino to piRNA clusters. Elife 11, e80067. 10.7554/eLife.80067.

50. Mohn, F., Sienski, G., Handler, D., and Brennecke, J. (2014). The rhino-deadlock-cutoff complex licenses noncanonical transcription of dual-strand piRNA clusters in Drosophila. Cell 157, 1364–1379. 10.1016/j.cell.2014.04.031.

51. Pane, A., Jiang, P., Zhao, D.Y., Singh, M., and Schupbach, T. (2011). The Cutoff protein regulates piRNA cluster expression and piRNA production in the Drosophila germline. EMBO J 30, 4601–4615. 10.1038/emboj.2011.334.

52. Chen, Y.A., Stuwe, E., Luo, Y., Ninova, M., Le Thomas, A., Rozhavskaya, E., Li, S., Vempati, S., Laver, J.D., Patel, D.J., et al. (2016). Cutoff Suppresses RNA Polymerase II Termination to Ensure Expression of piRNA Precursors. Mol Cell 63, 97–109. 10.1016/j.molcel.2016.05.010.

53. Zhang, Z., Wang, J., Schultz, N., Zhang, F., Parhad, S.S., Tu, S., Vreven, T., Zamore, P.D., Weng, Z.P., and Theurkauf, W.E. (2014). The HP1 Homolog Rhino Anchors a Nuclear Complex that Suppresses piRNA Precursor Splicing. Cell 157, 1353–1363. 10.1016/j.cell.2014.04.030.

54. Andersen, P.R., Tirian, L., Vunjak, M., and Brennecke, J. (2017). A heterochromatin-dependent transcription machinery drives piRNA expression. Nature 549, 54–59. 10.1038/nature23482.

55. Akkouche, A., Kneuss, E., Bornelöv, S., Renaud, Y., Eastwood, E.L., van Lopik, J., Gueguen, N., Jiang, M., Creixell, P., Maupetit-Mehouas, S., et al. (2024). A dual histone code specifies the binding of heterochromatin protein Rhino to a subset of piRNA source loci. bioRxiv 2024, 01.11.575256. 10.1101/2024.01.11.575256.

56. Sainsbury, S., Bernecky, C., and Cramer, P. (2015). Structural basis of transcription initiation by RNA polymerase II. Nat Rev Mol Cell Biol 16, 129–143. 10.1038/nrm3952.

57. Papai, G., Tripathi, M.K., Ruhlmann, C., Layer, J.H., Weil, P.A., and Schultz, P. (2010). TFIIA and the transactivator Rap1 cooperate to commit TFIID for transcription initiation. Nature 465, 956–960. 10.1038/nature09080.

58. Buratowski, S., Hahn, S., Guarente, L., and Sharp, P.A. (1989). Five intermediate complexes in transcription initiation by RNA polymerase II. Cell 56, 549–561. 10.1016/0092-8674(89)90578-3.

59. Malik, S., and Roeder, R.G. (2023). Regulation of the RNA polymerase II pre-initiation complex by its associated coactivators. Nat Rev Genet 24, 767–782. 10.1038/s41576-023-00630-9.

60. Thomas, M.C., and Chiang, C.M. (2006). The general transcription machinery and general cofactors. Crit Rev Biochem Mol Biol 41, 105–178. 10.1080/10409230600648736.

61. Naar, A.M., Lemon, B.D., and Tjian, R. (2001). Transcriptional coactivator complexes. Annu Rev Biochem 70, 475–501. 10.1146/annurev.biochem.70.1.475.

62. Bilgir, C., Dombecki, C.R., Chen, P.F., Villeneuve, A.M., and Nabeshima, K. (2013). Assembly of the Synaptonemal Complex Is a Highly Temperature-Sensitive Process That Is Supported by PGL-1 During Caenorhabditis elegans Meiosis. G3 (Bethesda) 3, 585-595. 10.1534/g3.112.005165.

63. Smothers, J.F., and Henikoff, S. (2000). The HP1 chromo shadow domain binds a consensus peptide pentamer. Curr Biol 10, 27–30. 10.1016/s0960-9822(99)00260-2.

64. Guven-Ozkan, T., Nishi, Y., Robertson, S.M., and Lin, R. (2008). Global transcriptional repression in C. elegans germline precursors by regulated sequestration of TAF-4. Cell 135, 149–160. 10.1016/j.cell.2008.07.040.

65. Feaver, W.J., Svejstrup, J.Q., Henry, N.L., and Kornberg, R.D. (1994). Relationship of CDK-activating kinase and RNA polymerase II CTD kinase TFIIH/TFIIK. Cell 79, 1103–1109. 10.1016/0092-8674(94)90040-x.

66. Shiekhattar, R., Mermelstein, F., Fisher, R.P., Drapkin, R., Dynlacht, B., Wessling, H.C., Morgan, D.O., and Reinberg, D. (1995). Cdk-Activating Kinase Complex Is a Component of Human Transcription Factor Tfiih. Nature 374, 283–287. 10.1038/374283a0.

67. Sun, Z.W., and Hampsey, M. (1995). Identification of the gene (SSU71/TFG1) encoding the largest subunit of transcription factor TFIIF as a suppressor of a TFIIB mutation in Saccharomyces cerevisiae. Proc Natl Acad Sci U S A 92, 3127–3131. 10.1073/pnas.92.8.3127.

68. Svejstrup, J.Q., Wang, Z., Feaver, W.J., Wu, X., Bushnell, D.A., Donahue, T.F., Friedberg, E.C., and Kornberg, R.D. (1995). Different forms of TFIIH for transcription and DNA repair: holo-TFIIH and a nucleotide excision repairosome. Cell 80, 21–28. 10.1016/0092-8674(95)90447-6.

69. Zaros, C., Briand, J.F., Boulard, Y., Labarre-Mariotte, S., Garcia-Lopez, M.C., Thuriaux, P., and Navarro, F. (2007). Functional organization of the Rpb5 subunit shared by the three yeast RNA polymerases. Nucleic Acids Res 35, 634–647. 10.1093/nar/gkl686.

70. Schilbach, S., Hantsche, M., Tegunov, D., Dienemann, C., Wigge, C., Urlaub, H., and Cramer, P. (2017). Structures of transcription pre-initiation complex with TFIIH and Mediator. Nature 551, 204–209. 10.1038/nature24282.

71. Nguyen, V.Q., Ranjan, A., Liu, S., Tang, X., Ling, Y.H., Wisniewski, J., Mizuguchi, G., Li, K.Y., Jou, V., Zheng, Q., et al. (2021). Spatiotemporal coordination of transcription preinitiation complex assembly in live cells. Mol Cell 81, 3560–3575. 10.1016/j.molcel.2021.07.022.

72. Kloc, A., Zaratiegui, M., Nora, E., and Martienssen, R. (2008). RNA interference guides histone modification during the S phase of chromosomal replication. Curr Biol 18, 490–495. 10.1016/j.cub.2008.03.016.

73. Reinhart, B.J., and Bartel, D.P. (2002). Small RNAs correspond to centromere heterochromatic repeats. Science 297, 1831. 10.1126/science.1077183.

74. Cam, H.P., Sugiyama, T., Chen, E.S., Chen, X., FitzGerald, P.C., and Grewal, S.I. (2005). Comprehensive analysis of heterochromatin- and RNAi-mediated epigenetic control of the fission yeast genome. Nat Genet 37, 809–819. 10.1038/ng1602.

75. Motamedi, M.R., Verdel, A., Colmenares, S.U., Gerber, S.A., Gygi, S.P., and Moazed, D. (2004). Two RNAi complexes, RITS and RDRC, physically interact and localize to noncoding centromeric RNAs. Cell 119, 789–802. 10.1016/j.cell.2004.11.034.

76. Schalch, T., Job, G., Noffsinger, V.J., Shanker, S., Kuscu, C., Joshua-Tor, L., and Partridge, J.F. (2009). High-affinity binding of Chp1 chromodomain to K9 methylated histone H3 is required to establish centromeric heterochromatin. Mol Cell 34, 36–46. 10.1016/j.molcel.2009.02.024.

77. Partridge, J.F., Scott, K.S., Bannister, A.J., Kouzarides, T., and Allshire, R.C. (2002). cis-acting DNA from fission yeast centromeres mediates histone H3 methylation and recruitment of silencing factors and cohesin to an ectopic site. Curr Biol 12, 1652–1660. 10.1016/s0960-9822(02)01177-6.

78. Law, J.A., Vashisht, A.A., Wohlschlegel, J.A., and Jacobsen, S.E. (2011). SHH1, a homeodomain protein required for DNA methylation, as well as RDR2, RDM4, and chromatin remodeling factors, associate with RNA polymerase IV. Plos Genet 7, e1002195. 10.1371/journal.pgen.1002195.

79. Law, J.A., and Jacobsen, S.E. (2010). Establishing, maintaining and modifying DNA methylation patterns in plants and animals. Nat Rev Genet 11, 204–220. 10.1038/nrg2719.

80. Haag, J.R., and Pikaard, C.S. (2011). Multisubunit RNA polymerases IV and V: purveyors of non-coding RNA for plant gene silencing. Nat Rev Mol Cell Biol 12, 483–492. 10.1038/nrm3152.

81. Haag, J.R., Ream, T.S., Marasco, M., Nicora, C.D., Norbeck, A.D., Pasa-Tolic, L., and Pikaard, C.S. (2012). In vitro transcription activities of Pol IV, Pol V, and RDR2 reveal coupling of Pol IV and RDR2 for dsRNA synthesis in plant RNA silencing. Mol Cell 48, 811-818. 10.1016/j.molcel.2012.09.027.

82. Law, J.A., Du, J., Hale, C.J., Feng, S., Krajewski, K., Palanca, A.M., Strahl, B.D., Patel, D.J., and Jacobsen, S.E. (2013). Polymerase IV occupancy at RNA-directed DNA methylation sites requires SHH1. Nature 498, 385–389. 10.1038/nature12178.

83. Le Thomas, A., Stuwe, E., Li, S., Du, J., Marinov, G., Rozhkov, N., Chen, Y.C., Luo, Y., Sachidanandam, R., Toth, K.F., et al. (2014). Transgenerationally inherited piRNAs trigger piRNA biogenesis by changing the chromatin of piRNA clusters and inducing precursor processing. Genes Dev 28, 1667–1680. 10.1101/gad.245514.114.

84. Towbin, B.D., Gonzalez-Aguilera, C., Sack, R., Gaidatzis, D., Kalck, V., Meister, P., Askjaer, P., and Gasser, S.M. (2012). Step-wise methylation of histone H3K9 positions heterochromatin at the nuclear periphery. Cell 150, 934–947. 10.1016/j.cell.2012.06.051.

85. Bessler, J.B., Andersen, E.C., and Villeneuve, A.M. (2010). Differential localization and independent acquisition of the H3K9me2 and H3K9me3 chromatin modifications in the Caenorhabditis elegans adult germ line. Plos Genet 6, e1000830. 10.1371/journal.pgen.1000830.

86. Chen, X., Xu, F., Zhu, C., Ji, J., Zhou, X., Feng, X., and Guang, S. (2014). Dual sgRNA-directed gene knockout using CRISPR/Cas9 technology in Caenorhabditis elegans. Sci Rep 4, 7581. 10.1038/srep07581.

87. Feng, G., Zhu, Z., Li, W.J., Lin, Q., Chai, Y., Dong, M.Q., and Ou, G. (2017). Hippo kinases maintain polarity during directional cell migration in Caenorhabditis elegans. EMBO J 36, 334–345. 10.15252/embj.201695734.

88. Kim, D., Paggi, J.M., Park, C., Bennett, C., and Salzberg, S.L. (2019). Graph-based genome alignment and genotyping with HISAT2 and HISAT-genotype. Nat Biotechnol 37, 907–915. 10.1038/s41587-019-0201-4.

