## Supplementary material for "The H3K27me3 reader UAD-2 recruits a TAF-12-containing transcription condensates to initiate piRNA expression within heterochromatic clusters": Supplemantary information

**Figure S1. TAF-12 is required for perinuclear localization of PRG-1 and piRNA biogenesis. a,** Putative top 10 interaction partners of UAD-2 identified by immunoprecipitation followed by mass spectrometry (IP-MS). **b,** Volcano plot comparing the transcriptomes of the indicated animals. Volcano plot showing $\log_2$ (fold change) of gene product-annotated transcript abundances in Auxin treatment vs control animals (x axis) and $-\log_{10}$ (P value) (y axis), calculated using an exact test in edgeR. The dashed line represents the significance threshold. The expression of *uad-2* mRNA is highlighted. **c,** Fluorescence images of representative mitotic and meiotic germ cells of young adult hermaphrodites expressing TAF-12::AID and GFP::PRG-1 (with or without auxin treatment). Scale bar, 10 μm. **d,** Western blotting analysis of the expression levels of GFP::3xFLAG::PRG-1 and β-actin with or without auxin treatment using anti-FLAG and anti-β-actin antibodies. **e,** Deep sequencing of total small RNAs in the indicated animals (with or without auxin treatment). Red dashed line indicates piRNAs.

**Figure S2. piRNA focus formation of the USTC complex depends on TAF-12. a-b,** Fluorescence images of representative embryos (**a**) and meiotic germ cells (**b**). TAF-12::GFP (green) colocalizes with the chromatin marker mCherry::HIS-58 (purple). **c,** Epidermal cells and intestinal cells of animals that express TAF-12::GFP. **d,** Fluorescence images of representative mitotic germ cells of the indicated animals with or without auxin treatment. Scale bar, 5 μm. **e-f,** Western blotting analysis of the expression levels of TAF-12::GFP::3xFLAG and β-actin in the indicated animals using anti-FLAG and anti-β-actin antibodies.

**Figure S3. UAD-2 interacts with TOFU-4. a,** Yeast two-hybrid assay to probe for protein-protein interactions between UAD-2, TAF-12, TOFU-4, SNPC-4, and PRDE-1. TOFU-5 is excluded because of its self-activation activity. On nonselective medium (left) all constructs allow growth equally, under selection (right) only strains expressing proteins that interact can grow. **b,** The amino acid sequence alignment of TAF-12 from different species was generated using MEGA and the ESPript3 server. Secondary structural elements of *C. elegans* TAF-12 is presented. **c,** Yeast two-hybrid assay to probe for protein-protein interactions between the UAD-2 chromodomain (UAD-2 chromo), the UAD-2 C-terminal domain (UAD-2 CTD), the TAF-12 histone-fold domain (TAF-12 HFD), and the TAF-12 C-terminal domain (TAF-12 CTD). On nonselective medium (left) all constructs allow growth equally, under selection (right) only strains expressing proteins that interact can grow.

**Figure S4. The TAF-12 C-terminal domain is essential for piRNA focus formation and piRNA production. a,** Schematic of the *taf-12(ust550/ΔCTD)* allele. The *taf-12(ust550/ΔCTD)* allele deletes 917 bp and does not induce frame shift in the open reading frame. **b,** Western blotting analysis of the expression levels of UAD-2::GFP::3xFLAG and β-actin in control and *taf-12(ΔCTD)* mutant worms using anti-FLAG and anti-β-actin antibodies. **c-e,** Fluorescence images of representative meiotic germ cells of the indicated animals expressing tagRFP::SYP-1 and TAF-12::GFP (**c**), TAF-12(*ΔCTD*)::GFP (**d**) and TAF-12(*ΔIDR*)::GFP (**e**). Scale bar, 5 μm. **f-g,** Scatter plots comparing the numbers of mature piRNA reads in the indicated animals.

**Figure S5. GTF-2F2, GTF-2H2C, MDT-8, and RPB-5 cooperate with TAF-12 to promote piRNA biogenesis. a,** List of *C. elegans* genes that encode major conserved transcription factors, including factors of the RNA Pol II, general transcription factors, the Mediator complex and the P-TEFb complex. **b-e,** Scatter plots comparing the numbers of mature piRNA reads in the indicated animals with or without auxin treatment. **f,** Fluorescence images of representative epidermal cells and intestinal cells of the indicated animals expressing indicated proteins. **g,** Yeast two-hybrid assay to probe for protein-protein interactions between UAD-2, TAF-12, MDT-8, GTF-2F2, GTF-2H2C, and RPB-5. On nonselective medium (left) all constructs allow growth equally, under selection (right) only strains expressing proteins that interact can grow.

**Table S1. The list of strains used in this study.**

**Table S2. The list of primers used for PCR amplification in this study.**

**Table S3. Sequences of the sgRNAs used in this study.**

**Figure S1**

**a**

Putative UAD-2-interacting partners (Top 10)

| WD score | Gene | Ortholog in *D. melanogaster* | Ortholog in *H. sapiens* | Brief description |
|---|---|---|---|---|
| 7024.68 | *uad-2* | - | - | piRNA biogenesis factor |
| 3761.63 | *taf-12* | Taf12-PA | TAF12 | Transcription factor |
| 649.6 | *F31D4.5* | IPIP-PD | PLEKHD1 | PH-like domain superfamily |
| 375.1 | *msp-76* | - | VAPA | Major sperm protein |
| 353.52 | *spk-1* | CG8565-PA | SRPK3 | SR protein kinase |
| 250.64 | *ZK673.2* | Ak3-PD | AK3 | Adenylate kinase activity |
| 215.69 | *T02D1.8* | - | - | Unknown function |
| 181.33 | *gyg-2* | Gyg-PE | GYG1 | Transferase activity |
| 135.15 | *ZK154.5* | - | - | Unknown function |
| 134.44 | *ssp-10* | - | - | Sperm specific family |

**b**

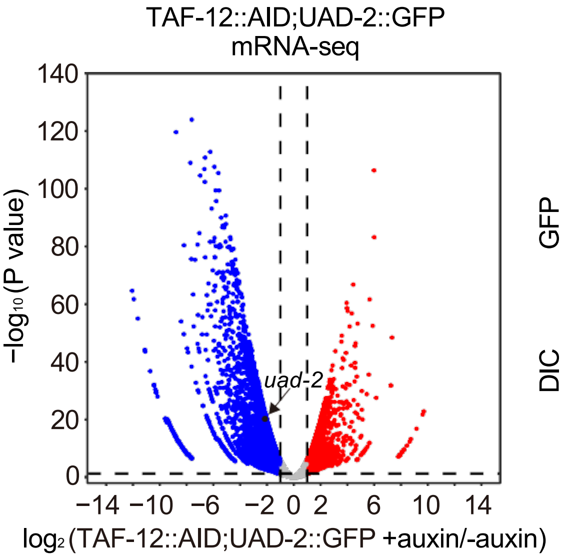


**c**

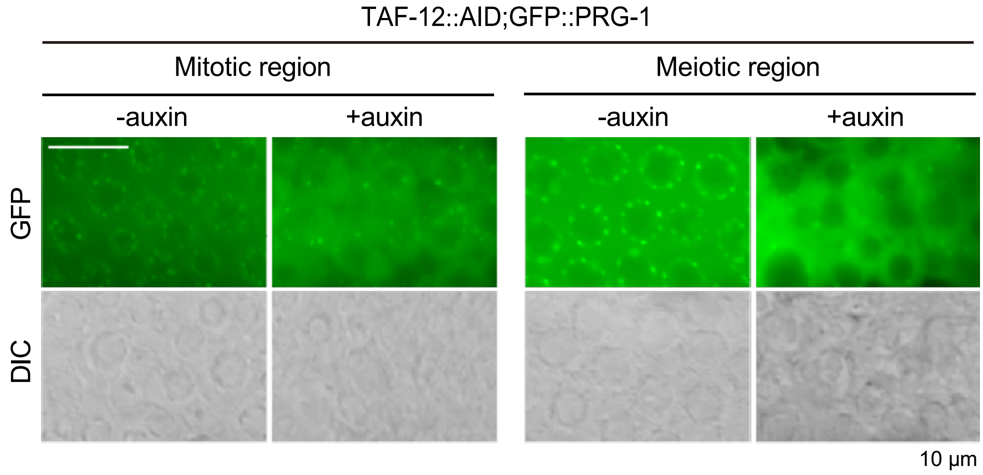


**d**

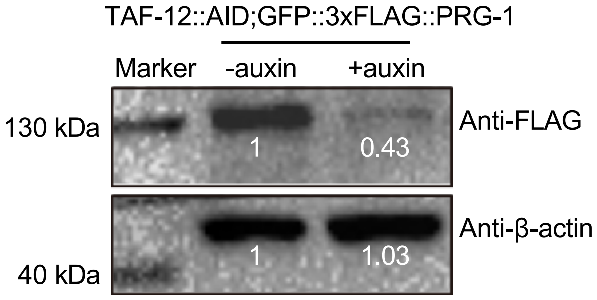


**e**

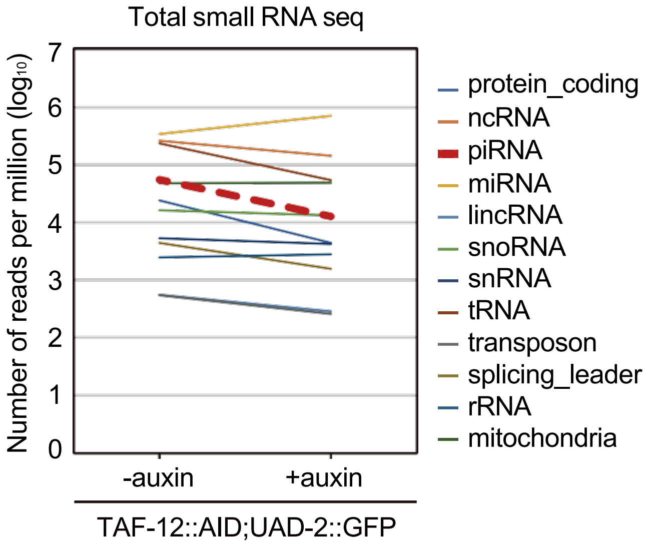

#### Figure S2

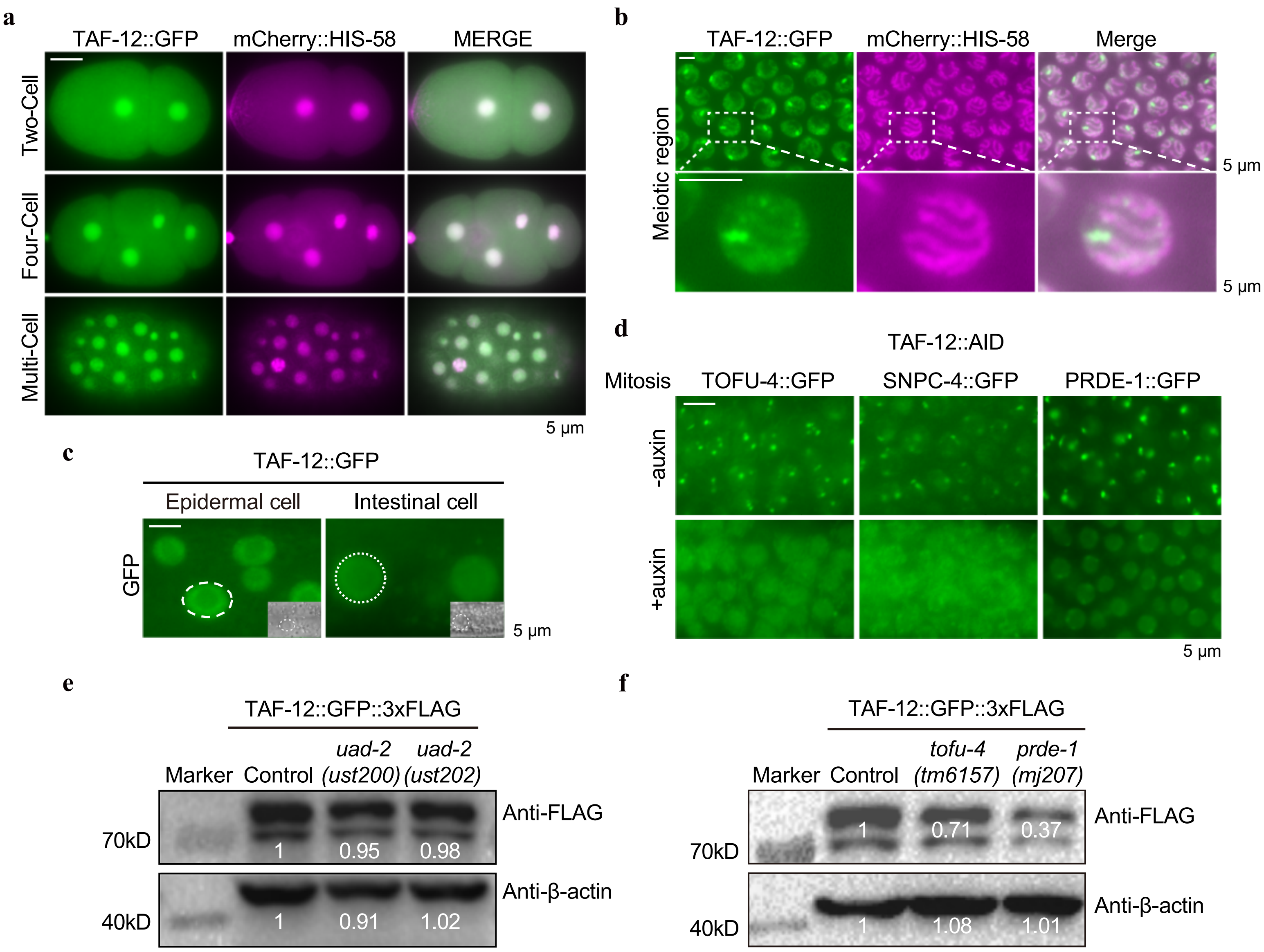
a
TAF-12::GFP
mCherry::HIS-58
MERGE
Two-Cell
Four-Cell
Multi-Cell
5 μm
b
TAF-12::GFP
mCherry::HIS-58
Merge
Meiotic region
5 μm
5 μm
c
TAF-12::GFP
Epidermal cell
Intestinal cell
GFP
5 μm
d
TAF-12::AID
Mitosis
TOFU-4::GFP
SNPC-4::GFP
PRDE-1::GFP
-auxin
+auxin
5 μm
e
TAF-12::GFP::3xFLAG
Marker
Control
uad-2 (ust200)
uad-2 (ust202)
70kD
1
0.95
0.98
Anti-FLAG
40kD
1
0.91
1.02
Anti-β-actin
f
TAF-12::GFP::3xFLAG
Marker
Control
tofu-4 (tm6157)
prde-1 (mj207)
70kD
1
0.71
0.37
Anti-FLAG
40kD
1
1.08
1.01
Anti-β-actin

### Figure S3

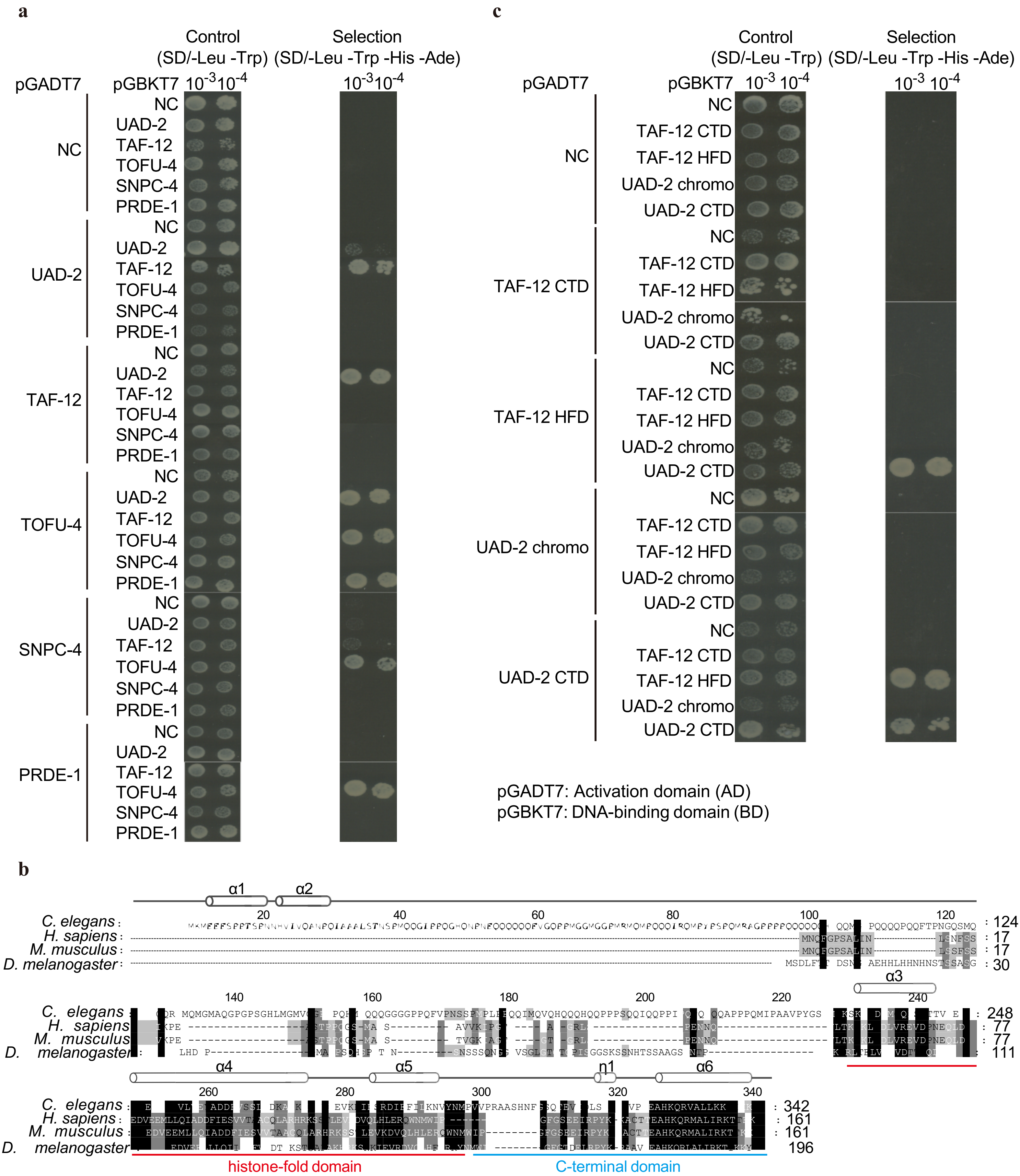
a
Control (SD/-Leu -Trp)
Selection (SD/-Leu -Trp -His -Ade)
pGADT7
pGBKT7
NC
UAD-2
TAF-12
TOFU-4
SNPC-4
PRDE-1
c
TAF-12 CTD
TAF-12 HFD
UAD-2 chromo
UAD-2 CTD
pGADT7: Activation domain (AD)
pGBKT7: DNA-binding domain (BD)
b
C. elegans
H. sapiens
M. musculus
D. melanogaster
histone-fold domain
C-terminal domain

### Figure S4

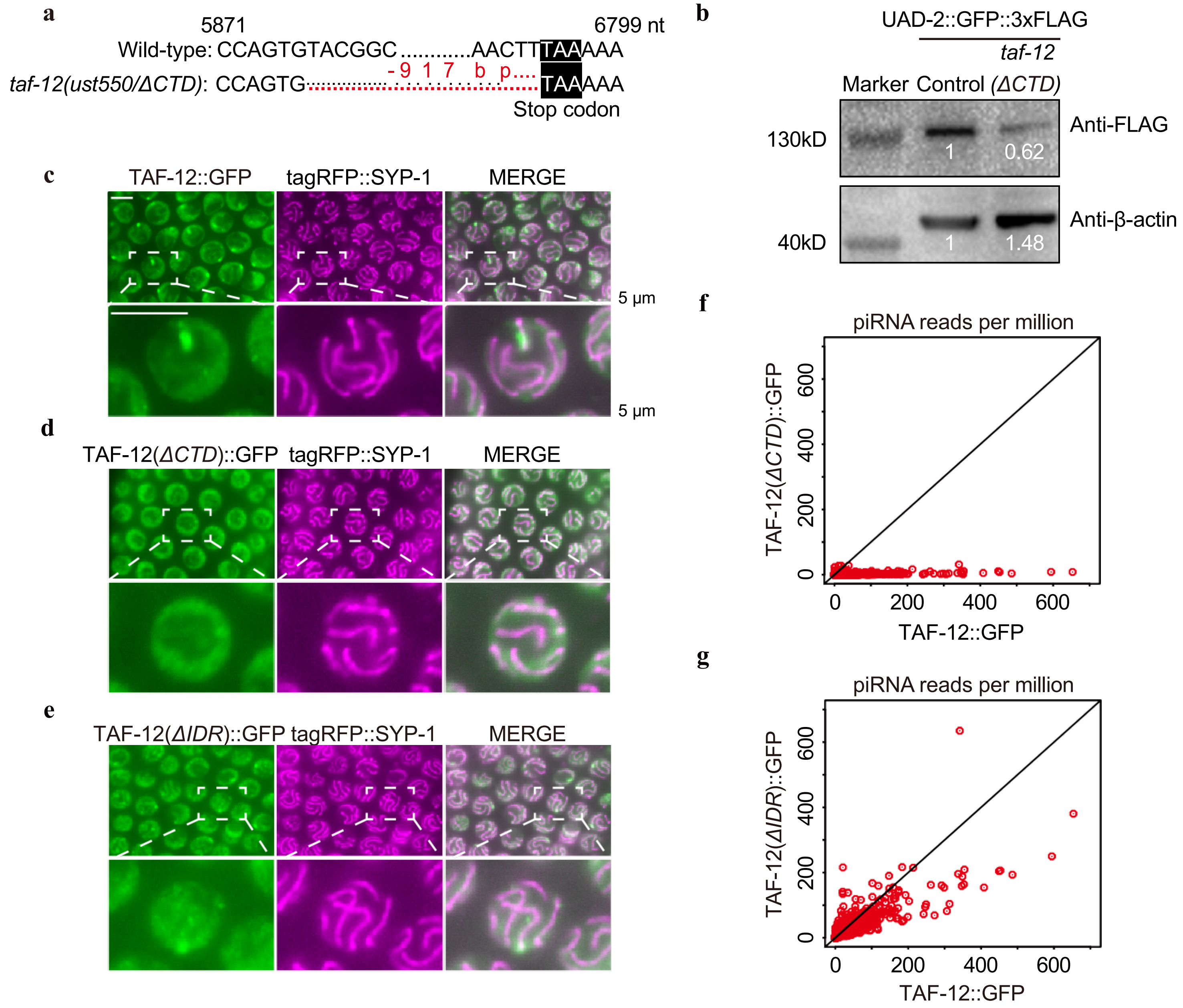
a
5871
6799 nt
Wild-type: CCAGTGTACGGC............AACTTTAAAAA
taf-12(ust550/ΔCTD): CCAGTG TAAAAA
-917 bp....
Stop codon
b
UAD-2::GFP::3xFLAG
taf-12
Marker Control (ΔCTD)
130kD
40kD
Anti-FLAG
1
0.62
Anti-β-actin
1
1.48
c
TAF-12::GFP
tagRFP::SYP-1
MERGE
5 μm
5 μm
d
TAF-12(ΔCTD)::GFP
tagRFP::SYP-1
MERGE
e
TAF-12(ΔIDR)::GFP
tagRFP::SYP-1
MERGE
f
piRNA reads per million
TAF-12(ΔCTD)::GFP
TAF-12::GFP
0
200
400
600
g
piRNA reads per million
TAF-12(ΔIDR)::GFP
TAF-12::GFP
0
200
400
600

#### Figure S5

**a**

***C. elegans* genes that encode major conserved transcription factors**

| Complex | *C. elegans* genes |
|---|---|
| **RNA Pol II** | *ama-1, rpb-2, rpb-3, rpb-4, rpb-5, rpb-6, rpb-7, rpb-8, rpb-9, rpb-10, rpb-11, rpb-12* |
| **TFIIA** | *pqn-51, gtf-2A2* |
| **TFIIB** | *ttb-1* |
| **TFIID** | *taf-1, taf-2, taf-3, taf-4, taf-5, taf-6.1, taf-6.2, taf-7.1, taf-7.2, taf-8, taf-9, taf-10, taf-11.1, taf-11.2, taf-12, taf-13* |
| **TFIIE** | *gtf-2E2* |
| **TFIIF** | *gtf-2F2* |
| **TFIIH** | *xpb-1, xpd-1, gtf-2H2C, gtf-2H3, cdk-7, cyh-1, mnat-1* |
| **Mediator** | *mdt-1.1, mdt-1.2, mdt-4, mdt-6, mdt-7, mdt-8, mdt-10, mdt-11, mdt-12, mdt-13, mdt-14, mdt-15, mdt-17, mdt-18, mdt-19, mdt-20, mdt-21, mdt-22, mdt-23, mdt-27, mdt-29, mdt-31, cdk-8, cic-1* |
| **P-TEFb** | *cit-1.1, cit-1.2* |

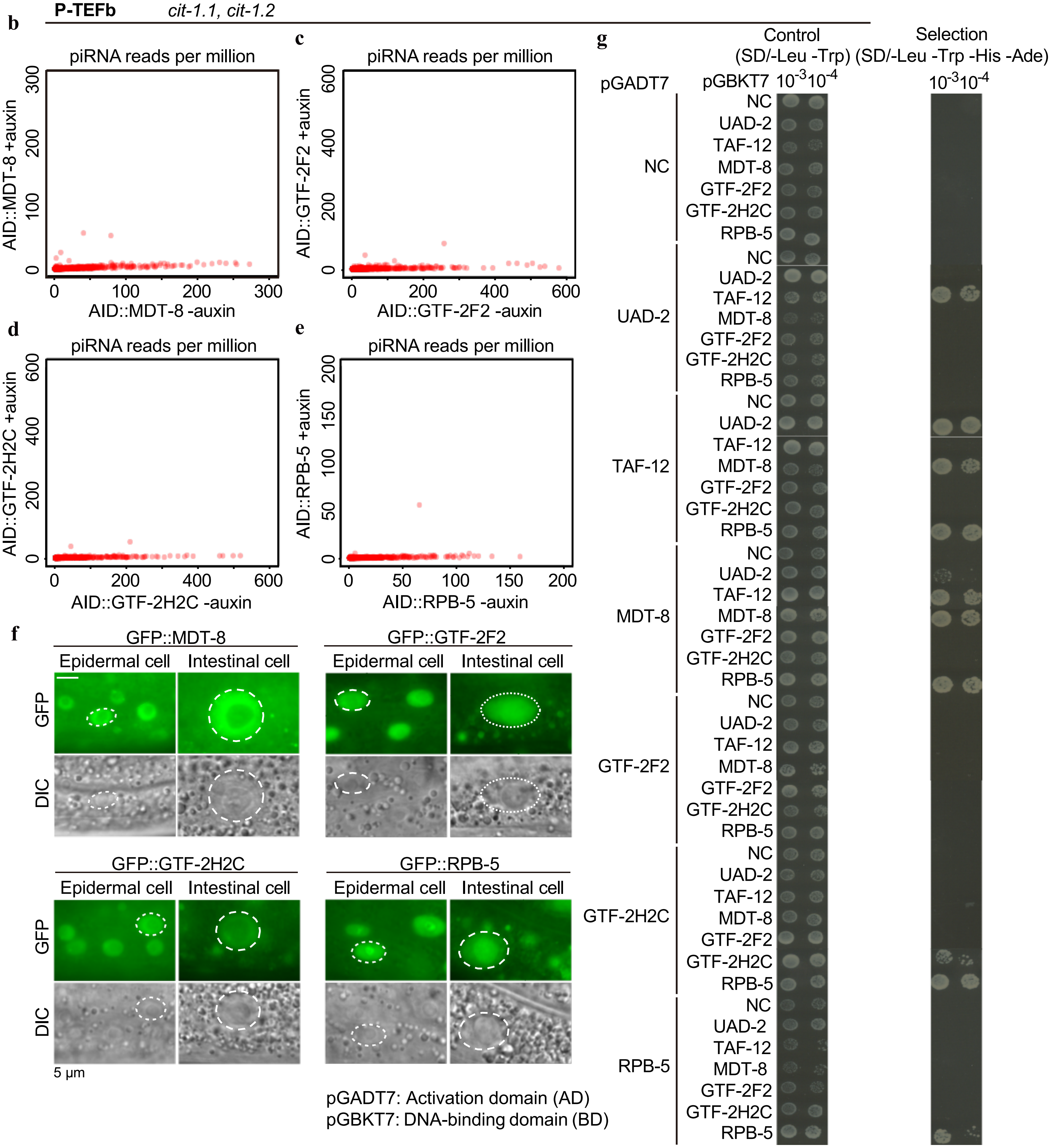

**Table S1. The list of strains used in this study.**

| Strain | Allele |
|---|---|
| N2 | / |
| SHG1093 | *uad-2(ust654[uad-2::gfp::3xflag]) I* |
| SHG1745 | *taf-12(ust652[taf-12::gfp::3xflag]) III* |
| SHG1746 | *taf-12(ust653[taf-12::AID]) III; ieSi38[sun-1p::TIR1::mRuby::sun-1_3'UTR + Cbr-unc-119(+)] IV; uad-2(ust654[uad-2::gfp::3xflag]) I* |
| SHG1795 | *taf-12(ust652[taf-12::gfp::3xflag]) III; ltIs37[pie-1p::mCherry::his-58 + unc-119(+)] IV* |
| SHG1874 | *taf-12(ust653[taf-12::AID]) III; ieSi38[sun-1p::TIR1::mRuby::sun-1_3'UTR + Cbr-unc-119(+)] IV; prg-1(ust643[gfp::3xflag::prg-1]) I* |
| SHG1876 | *taf-12(ust653[taf-12::AID]) III; ieSi38[sun-1p::TIR1::mRuby::sun-1_3'UTR + Cbr-unc-119(+)] IV; ustIS028(tofu-4p::tofu-4::gfp::3xflag::tofu-4_3'UTR) II* |
| SHG1901 | *taf-12(ust653[taf-12::AID]) III; ieSi38[sun-1p::TIR1::mRuby::sun-1_3'UTR + Cbr-unc-119(+)] IV; vrIs87[pie-1p::snpc-4::gfp::snpc-4_3'UTR + Cbr-unc-119(+)]* |
| SHG1902 | *rpb-5(ust655[3xflag::bfp::AID::rpb-5]) I; uad-2(ust654[uad-2::gfp::3xflag]) I* |
| SHG1903 | *gtf-2f2(ust656[3xflag::bfp::AID::GTF-2F2]) V; uad-2(ust654[uad-2::gfp::3xflag]) I* |
| SHG2095 | *gtf-2h2c(ust657[3xflag::bfp::AID::gtf-2h2c]) III; uad-2(ust654[uad-2::gfp::3xflag]) I* |
| SHG2096 | *mdt-8(ust658[3xflag::bfp::AID::mdt-8]) II; uad-2(ust654[uad-2::gfp::3xflag]) I* |
| SHG2181 | *gtf-2h2c(ust384[3xflag::gfp::gtf-2h2c]) III; mjSi74[mex-5p::wormCherry::prde-1::par-5_3'UTR] I* |
| SHG2382 | *mdt-8(ust425[3xflag::gfp::mdt-8]) II; mjSi74[mex-5p::wormCherry::prde-1::par-5_3'UTR] I* |
| SHG2520 | *uad-2(ust200) I; taf-12(ust652[taf-12::gfp::3xflag]) III; ustIS325(mex-5p::3xha::tagRFP::SYP-1::tbb-2_3'UTR) I* |
| SHG2542 | *taf-12(ust660[taf-12(del IDR)::gfp::3xflag]) III; ustIS325(mex-5p::3xha::tagRFP::SYP-1::tbb-2_3'UTR) I* |
| SHG2549 | *uad-2(ust654[uad-2::gfp::3xflag]) I; taf-12(ust485[taf-12::tagRFP::3xha]) III* |
| SHG2675 | *taf-12(ust465[taf-12(del C-terminal domain)::gfp::3xflag]) III; ustIS325(mex-5p::3xha::tagRFP::SYP-1::tbb-2_3'UTR) I* |
| SHG2707 | *taf-12(ust652[taf-12::gfp::3xflag]) III; ustIS325(mex-5p::3xha::tagRFP::SYP-1::tbb-2_3'UTR) I* |
| SHG2728 | *prde-1(mj207) V; taf-12(ust652[taf-12::gfp::3xflag]) III; ustIS325(mex-5p::3xha::tagRFP::SYP-1::tbb-2_3'UTR) I* |
| SHG2772 | *taf-12(ust550) III* |
| SHG2841 | *taf-12(ust550) III; snpc-4(ust414[snpc-4::gfp::3xflag]) I* |
| SHG2850 | *taf-12(ust550) III; prde-1(ust393[prde-1::gfp::3xflag]) V* |
| SHG2865 | *taf-12(ust550) III; tofu-4(ustIS248[tofu-4::gfp::3xflag]) I* |
| SHG2895 | *taf-12(ust653[taf-12::AID]) III; ieSi38[sun-1p::TIR1::mRuby::sun-1_3'UTR + Cbr-unc-119(+)] IV; prde-1(ust393[prde-1::gfp::3xflag]) V* |
| SHG3004 | *taf-12(ust550) III; uad-2(ust654[uad-2::gfp::3xflag]) I* |
| SHG3012 | *taf-12(ust652[taf-12::gfp::3xflag]) III; mjSi74[mex-5p::wormCherry::prde-1::par-5_3'UTR] I* |
| SHG3027 | *tofu-4(tm6157) I; taf-12(ust652[taf-12::gfp::3xflag]) III; ustIS325(mex-5p::3xha::tagRFP::SYP-1::tbb-2_3'UTR) I* |
| SHG3028 | *gtf-2f2(ust611[3xflag::gfp::gtf-2f2]) V; mjSi74[mex-5p::wormCherry::prde-1::par-5_3'UTR] I* |
| SHG3067 | *rpb-5(ust661[3xflag::gfp::rpb-5]) I; mjSi74[mex-5p::wormCherry::prde-1::par-5_3'UTR] I* |
| SHG3148 | *taf-12(ust660[taf-12(del IDR)::gfp::3xflag]) III; mjSi74[mex-5p::wormCherry::prde-1::par-5_3'UTR] I* |
| SHG3177 | *taf-12(ust550) III; uad-2(ust654[uad-2::gfp::3xflag]) I; ustSi336(mex-5p::taf-12::tagRFP::3xha::tbb-2_3'UTR) II* |
| SHG3208 | *taf-12(ust653[taf-12::AID]) III; ieSi38[sun-1p::TIR1::mRuby::sun-1_3'UTR + Cbr-unc-119(+)] IV; ustSi336(mex-5p::taf-12::tagRFP::3xha::tbb-2_3'UTR) II* |

**Table S2. The list of primers used for PCR amplification in this study.**

| Primer | sequence |
|---|---|
| *gfp::3xflag* F | GGAGGTGGAGGTGGAGCTAT |
| *gfp::3xflag* R | cttgtcatcgtcatccttgtaatcga |
| *tagRFP::3xha* F | GGAGGTGGAGGTGGAGCTATG |
| *tagRFP::3xha* R | GTAATCTGGAACATCGTATGGGTAAGCGTAATCTGGAACATCGTATGGGTAGTTGAGCTTGTGCCCG |
| *taf-12::gfp::3xflag/taf-12::AID* Left arm F | gggtaacgccagCACGTGtggctcctgaaaaaataatgaaatt |
| *taf-12::gfp::3xflag/taf-12::AID* Left arm R | ATAGCTCCACCTCCACCTCCAAGTTTCTTGATTTGTTTCTTG |
| *taf-12::gfp::3xflag* Right arm F | acaaggatgacgatgacaagTAAaaattaatttttttttgttttaattttaattttc |
| *taf-12::gfp::3xflag* Right arm R | cagcggataacaatttcacacatcaacaacaatatcggt |
| *taf-12::AID* Right arm F | GGAGGTGGAGGTGGAGCTatgcctaaagatccagccaaacctc |
| *taf-12::AID* Right arm R | cttcacgaacgccgccg |
| *3xflag::gfp* F | gactacaaagaccatgacgg |
| *3xflag::gfp* R | AAGGAGGTGGAGGTGGAGCT |
| *AID::rpb-5/gfp::RPB-5* Left arm F | gggtaacgccagCACGTGtggcaagggtacatcagcggga |
| *AID::rpb-5/gfp::RPB-5* Left arm R | ccgtcatggtctttgtagtcCTCGTCGTCAGCCATtatta |
| *AID::rpb-5/gfp::RPB-5* Right arm F | cagcggataacaatttcacaccacatctgtgctcatttcg |
| *AID::rpb-5/gfp::RPB-5* Right arm R | cagcggataacaatttcacaccacatctgtgctcatttcg |
| *AID::mdt-8/gfp::mdt-8* Left arm F | gggtaacgccagCACGTGtgggcactgtgccaaacgcaca |
| *AID::mdt-8/gfp::mdt-8* Left arm R | ccgtcatggtctttgtagtcATTTGGATACTCCATctgaa |
| *AID::mdt-8/gfp::mdt-8* Right arm F | GGAGGTGGAGGTGGAGCTATGGAGTATCCAAATCCACC |
| *AID::mdt-8/gfp::mdt-8* Right arm R | cagcggataacaatttcacattcgtcattttgggaagagg |
| *AID::gtf-2h2c/gfp::gtf-2h2c* Left arm F | gggtaacgccagCACGTGtgtcgagcatcatatgcaggaa |
| *AID::gtf-2h2c/gfp::gtf-2h2c* Left arm R | ccgtcatggtctttgtagtcCTCATCATCATCCATattta |
| *AID::gtf-2h2c/gfp::gtf-2h2c* Right arm F | AAGGAGGTGGAGGTGGAGCTATGGATGATGATGAGCAGAA |
| *AID::gtf-2h2c/gfp::gtf-2h2c* Right arm R | cagcggataacaatttcacaGAACATTTCAGCAGACAGTC |
| *AID::gtf-2f2/gfp::gtf-2f2* Left arm F | gggtaacgccagCACGTGtggcaattttcgcactgaaatc |
| *AID::gtf-2f2/gfp::gtf-2f2* Left arm R | ccgtcatggtctttgtagtcCTTCGTTGGGCTCATtttcc |
| *AID::gtf-2f2/gfp::gtf-2f2* Right arm F | AAGGAGGTGGAGGTGGAGCTATGAGCCCAACGAAGCGATA |
| *AID::gtf-2f2/gfp::gtf-2f2* Right arm R | cagcggataacaatttcacaTAATTCGTCCCTCAATTGCC |
| *3xflag::BFP::AID* linker F | gactacaaagaccatgacgg |
| *3xflag::BFP::AID* linker R | AGCTCCACCTCCACCTCCct |

**Table S3. Sequences of the sgRNAs used in this study.**

| sgRNA targets | sgRNA sequences |
|---|---|
| *taf-12* 3' sg#1 | GAAGGAAAAGTTTGTTCCAA |
| *taf-12* 3' sg#2 | GCACAATTTTGGGTCTCAAA |
| *taf-12* 3' sg#3 | AACGGAAGCACACAAGCAAC |
| *taf-12* 3' sg#4 | AGTGTTGGAGGTATTACTCA |
| *taf-12* 3' sg#5 | TACCTCCAACACTTCAGATT |
| *gtf-2f2* 5' sg#1 | AGAGCTCTGGGATGCCAACG |
| *gtf-2f2* 5' sg#2 | GAATGACGTCGACTGTGAGC |
| *gtf-2f2 5'* sg#3 | GTTGTTGGAAAATTGCAGAT |
| *gtf-2h2c 5'* sg#1 | TCAAGGTGAAAAGTATTACT |
| *gtf-2h2c 5'* sg#2 | ATGAGCAGAAGGGTTACACC |
| *gtf-2h2c 5'* sg#3 | TAATAAAAGCCACTTCCAGA |
| *mdt-8 5'* sg#1 | TCACTCTTTTGATCATCATA |
| *mdt-8 5'* sg#2 | ATCTGATTGCTGGACACGGA |
| *mdt-8 5'* sg#3 | ATAGACATCTTAAAATCTTA |
| *rpb-5 5'* sg#1 | ACGGATTCTCCACAATCTAT |
| *rpb-5 5'* sg#2 | AGATGAATTAGATCAACCAT |
| *rpb-5 5'* sg#3 | TATTTTTCAACCTTTAATGT |
| *taf-12(del IDR*) sg#1 | TTCTTCATACTTTGCTGCTG |
| *taf-12* (*del IDR*) sg#2 | TTCGACCAATCAGGTAGATT |
| *taf-12* (*del IDR*) sg#3 | ACTTGCATAATTTGTTGAGG |
| *taf-12* (*del IDR*) sg#4 | TGAATTTGTTGAGATGGTGG |
